# Filament structure and subcellular organization of the bacterial intermediate filament-like protein crescentin

**DOI:** 10.1101/2023.06.04.543601

**Authors:** Yue Liu, Fusinita van den Ent, Jan Löwe

**Author notes:** Correspondence: Jan Löwe, MRC Laboratory of Molecular Biology, Cambridge Biomedical Campus, Francis Crick Avenue, Cambridge CB2 0QH, UK. Present address: Department of Microbiology and State Key Laboratory for Diagnosis and Treatment of Infectious Diseases of the First Affiliated Hospital, Zhejiang University School of Medicine, Hangzhou, Zhejiang 310058, China (Yue Liu). **CLASSIFICATION:** Biological Sciences, Biophysics and Computational Biology.

## Abstract

The coiled coil protein crescentin is required for the crescent shape of the freshwater bacterium *Caulobacter crescentus* (*vibrioides*). Crescentin forms a filamentous structure on the inner, concave side of the curved cells. It shares features with eukaryotic intermediate filament (IF) proteins, such as its ability to form filaments *in vitro*, the protein’s length, sequence comparisons and the presence of a coiled coil discontinuity called the “stutter”. Here, we have used electron cryomicroscopy (cryo-EM) to determine the structure of the full-length protein and its filament, exploiting a crescentin-specific nanobody. The filament is formed by two strands, related by two-fold symmetry that each consist of two dimers, resulting in an octameric assembly. Crescentin subunits form longitudinal contacts head-to-head and tail-to-tail, making the entire filament non-polar. Using *in vivo* site-directed cysteine crosslinking we demonstrated that contacts observed in the *in vitro* filament structure exist in cells. Electron cryotomography (cryo-ET) of cells expressing crescentin showed filaments on the concave side of the curved cells, close to the inner membrane, where they form a band. Comparison of our crescentin filament structure with current models of IF proteins and their filaments revealed similar coiled coil dimer formation as well as an absence of overall polarity. IF proteins form head-to-tail longitudinal contacts in contrast to crescentin and hence several inter-dimer contacts in IFs have no equivalents in crescentin filaments. Our work supports the idea that intermediate filament-like proteins achieve their shared polymerization and mechanical properties through a variety of filament architectures.

**SIGNIFICANCE STATEMENT:** Crescentin is a coiled coil protein that is required for the crescent cell shape of bacteria such as *Caulobacter crescentus*. Crescentin shares biochemical and cytoskeletal properties with intermediate filament (IF) proteins, which form the third major class of cytoskeletal proteins in eukaryotes. To better understand the relationship between crescentin and IF proteins, and the filaments they form, we have determined the three-dimensional structure of crescentin filaments by cryo-EM. This revealed the full-length structure of the parallel coiled coil dimer of crescentin and how dimers come together laterally and longitudinally, to form a non-polar, octameric filament. Differences in filament architecture highlight the versatility of intermediate filament-like proteins across the tree of life.

## INTRODUCTION

Fitness of bacteria in the environment is often dependent on their shape. It is therefore that most bacteria tightly control their shape, producing spheres (cocci), rods (bacilli), helices, filaments and irregular, appended as well as crescent shapes [1]. In many cases, shape is dependent on the stress-bearing cell wall. Hence shape determination is dependent on the ability to regulate cell wall synthesis and remodelling during vegetative growth, as well as during cell division [2].

*Caulobacter crescentus* (*vibrioides*) is crescent shaped. The Gram-negative freshwater bacterium has been studied extensively, and not only because its cells undergo a developmental transition from swarmer to stalked cells. The larger stalked cells are able to attach to surfaces via the stalk, while the swarmer cells are flagellated and motile. Only stalked cells replicate DNA and divide. Their cell division is asymmetric and leads to one swarmer cell and one stalked cell, closing the cell cycle [3]. Both cell types are crescent-shaped. It is thought that the crescent shape is advantageous for cell mobility in their aqueous environment, but it has recently been suggested that the crescent shape enables pili on the swarmer cells to attach to surfaces for biofilm formation [4].

Whatever the biological reason for its crescent shape, in *Caulobacter* it is dependent on the function of the protein crescentin (CreS) [5]. Immunofluorescence and fluorescent protein-tagging using a merodiploid strain (because CreS-GFP fusions were non-functional) showed crescentin to form a single filamentous structure on the inner (concave) surface of the cell crescent [5–7]. The proximity of the crescentin structure to the cell membrane was lost during treatment with the MreB inhibitor A22 or the cell wall inhibitors mecillinam and phosphomycin [6]. Based on the cellular localization, the cytoskeletal properties of crescentin and modelling it has been proposed that its mode of action to create cell shape is mechanical [6]. The finding that MreB is involved [7] might indicate that crescentin filaments reduce cell wall synthesis on one side of the cell, leading to the crescent shape. In line with these ideas, it was found that the heterologous expression of crescentin in *E. coli* leads to curved cells [6], suggesting a conserved or very simple mechanism of cell wall synthesis inhibition.

Crescentin levels in cells do not vary much during the cell cycle [7]. Crescentin filaments in cells are not dynamic and free molecules are added to the filamentous cellular structure rapidly, indicating strong cooperativity of assembly [7, 8]. Initial assembly starts with a cell-long thin structure that thickens as the cell progresses through the cell cycle. Re-arrangements of the crescentin structure occur during cell division, although it is not known how those are regulated or facilitated [7].

The discovery of crescentin as a regulator of cell shape attracted a lot of attention also because of a number of similarities to eukaryotic intermediate filament proteins (IFs) [5, 9]. IFs are the third major class of intracellular filaments in eukaryotes, in addition to microfilaments (actin) and microtubules (tubulin). IFs can be categorized into five major families and include two types of cytoplasmic keratins, vimentin and neurofilament, as well as nuclear lamins [10]. IFs have a well-defined domain architecture, with a central α-helical “rod” domain driving the formation of parallel coiled coil dimers. The rod domain, which has been sub-divided into three coiled coil segments (coil 1A, 1B and 2) and two inter-connecting linkers (L1 and L12), is complemented primarily by disordered head and tail domains (in contrast, nuclear lamins and invertebrate cytoplasmic IFs have folded Ig-fold tail domains). Cytoplasmic IFs have rod domains of about 308 residues, nuclear lamins of about 350 residues. All IF sequences have a “stutter” in the third rod segment that deviates from the regular coiled coil heptad repeat pattern. Despite many years of research there is still no complete structural description of any IF filament [11]. They are known to form tetramers through inter-dimer interactions mediated by their N-terminal regions of the rod domain (“A11” inter-dimer contact). Filaments are thought to contain tetramers as building blocks, at least for the cytoplasmic IFs, and three other types of inter-dimer contacts (“A22”, “A12”, and “ACN”) then lead to higher-order structures such as the prototypical 10 nm-thick mature fibres of many IF proteins [12].

Crescentin contains the heptad repeats needed to form an extended coiled coil dimer, and also a stutter in the C-terminal half. The rod domain of crescentin is longer than IF proteins, and it was demonstrated early on that crescentin forms very stable filaments *in vitro* [5]. Filament formation *in vitro* and *in vivo* showed that the domain organization of crescentin is important for its polymerization and cell shape maintenance functions [9]. In particular, membrane attachment was shown to depend on the first 27 amino acids [6, 9]. Removal of the stutter also interfered with crescentin’s ability to curve cells [9].

Given the enigmatic relationship between crescentin and IFs, our lack of understanding how crescentin filaments form in cells and how they control cell shape, we set out to understand the crescentin filament structure. We employed electron cryomicroscopy (cryo-EM) on crescentin filaments bound by a newly developed megabody [13] to determine their structure, which revealed an “octameric” twisted and double-stranded filament without polarity. We used *in vivo* site-specific cysteine crosslinking to demonstrate that many, if not all features of the *in vitro* filament structure exist in crescentin filaments in cells. Electron cryotomography (cryo-ET) of *Caulobacter* cells expressing crescentin showed the filaments on the inner, concave side of cells, close to the inner membrane where they form a single, wide band. Comparison with current models of IF proteins revealed significant differences between their filament architectures.

## RESULTS

### Development of a megabody for CreS structure determination

In line with previously reported assembly properties of crescentin (CreS) and of many eukaryotic IF proteins [5, 9, 12, 14–16], polymerization of purified, un-tagged *Caulobacter crescentus (vibrioides)* CreS upon a decrease of pH *in vitro* led to CreS aggregates, bundles, or irregular filaments that exhibited structural polymorphism. In some cases, cryo-EM micrographs of wild-type (wt) CreS, namely CreS_wt_, polymerized at pH 7.0 showed unbundled filaments with a width of ∼9 nm (**Fig. S1A**). Two-dimensional (2D) class averages revealed two intertwined strands with a regular spacing of ∼57 nm (distance between helical crossovers) (**Fig. S1B**). Nevertheless, the smooth appearance of these filaments prevented determination of the register of individual subunits along the filament axis.

To facilitate structure determination of CreS, we raised nanobodies (NBs) against purified CreS_wt_ and selected NBs that bind to CreS_wt_ polymers formed at low pH conditions (**Materials and Methods**). Coiled coil prediction [17] and three-dimensional (3D) structure prediction by AlphaFold 2 (AF2) [18] **(Fig. S1C)** indicated that CreS contains a central, coiled coil “rod” domain (residues 80-444), flanked by an N-terminal segment (residues 1-79, although somewhat shorter in AF2 predictions, **Fig. S1C**) and a C-terminal segment (residues 445-457) (**Fig. 1A**). Given the elongated nature of the CreS coiled coil dimer, we engineered an NB derivative, megabody (MB), to act as a bulky and rigid marker bound to CreS. For each NB investigated, a MB was generated by grafting the NB onto a circular permutant of the scaffold protein *E. coli* YgjK, as previously described [13]. Among the MBs, megabody 13 (MB13), a derivative of nanobody 13 (NB13) binds to both CreS_wt_ and CreS_sat_ at pH 8. CreS_sat_ contains a stretch of three amino acids (Ser, Ala and Thr; SAT), inserted before residue 406, to remove the “stutter” that locally disrupts the continuity of the heptad-repeat coiled coil [9, 19] (**Figs. 1B-C and S1D-E**). Cryo-EM analysis revealed that for both versions of CreS, purified CreS-MB13 complex at pH 8 polymerizes into regular, unbundled filaments upon lowering the pH to 6.5 in the presence of detergents, such as CHAPS (**Fig. 1D**). These filaments showed a regular spacing of ∼57 nm between neighbouring “nodes”, where a node is a segment of the CreS filament decorated with multiple MB13 molecules (**Figs. 1E and S3A**).

**Figure 1.**
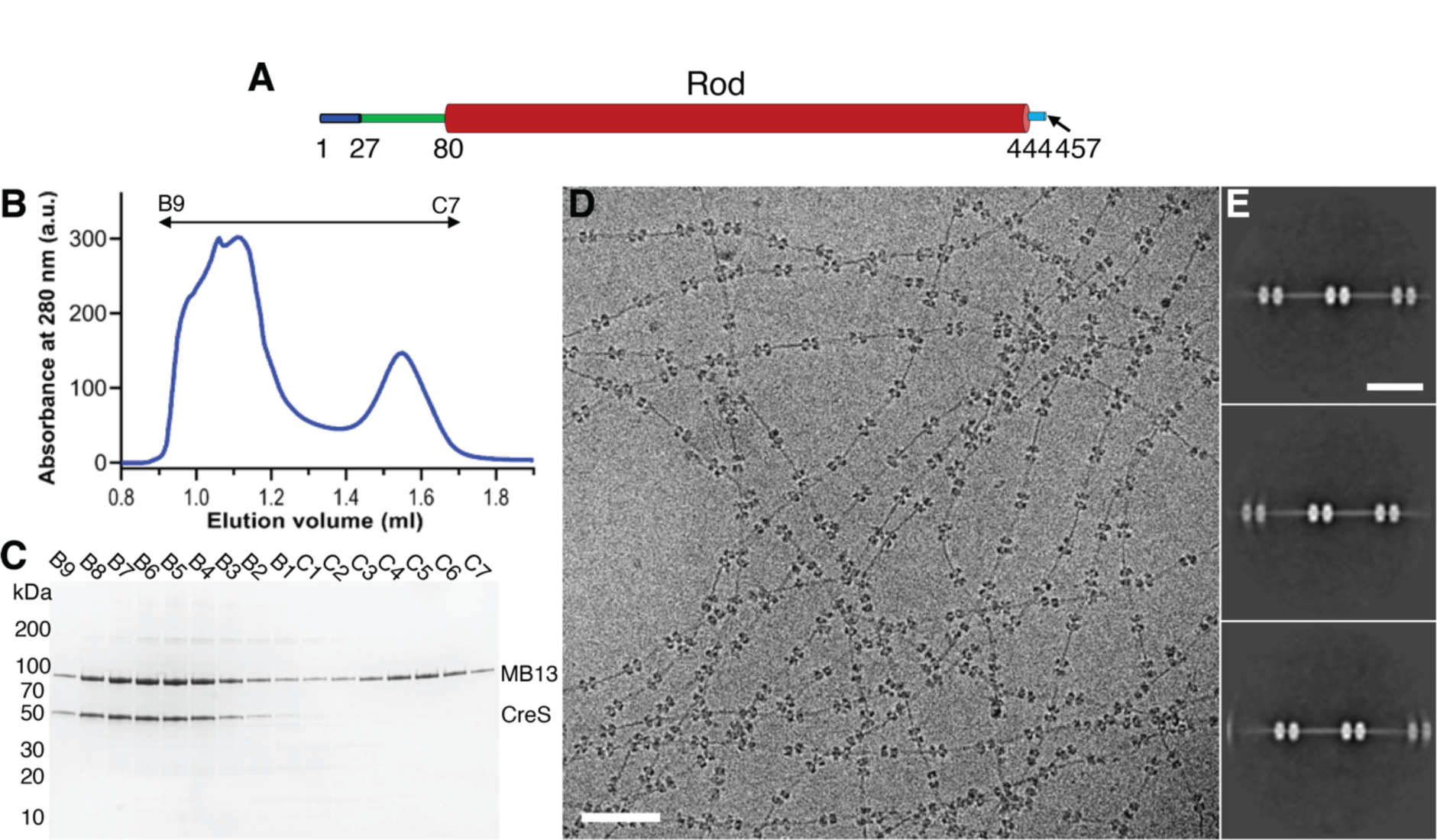
Production and characterization of a CreS-specific megabody (MB) for structure determination of CreS filaments. **(A)** Domain organization of full length CreS. The rod domain was defined based on coiled coil prediction [17] and structure prediction using AlphaFold 2 [18] (see **Fig. S1C**). **(B**) Size exclusion chromatography (SEC) profile of CreS_sat_ (a mutant version that has a three amino acid insertion [Ser, Ala and Thr, SAT] before position 406), preincubated with MB13 at 1:4 molar ratio at pH 8. See **Fig. S1D-E** for SEC profile of CreS_wt_ mixed with MB13. (**C**) SDS-PAGE analysis of SEC fractions. Fractions B9 to B5 were pooled and used for EM studies. (**D**) A typical cryo-EM image showing single CreS filaments in complex with MB13 that are formed at pH 6.5. (**D**) Reference-free 2D classification of these filaments reveals a regular spacing of ∼60 nm between neighbouring “nodes” along the filament. The nodes are caused by MB binding.

For cryo-EM structure determination of CreS, we treated nodes as single particles, using a box size of ∼420 Å (henceforth referred to as “small box”), to obtain well-resolved reconstructions of CreS bound with NB13 molecules (**Materials and Methods, Table S3, Fig. S2**). We next extended the box size to ∼960 Å (referred to as “large box”) for the visualization of a more complete CreS structure, which included regions that are distant from the NB13 binding site, which was mapped to a segment near the C-terminus of CreS (**Materials and Methods, Fig. 2A, Table S3, Fig. S2**). The best small box and large box reconstructions were of CreS_sat_ complexed with NB13, determined at nominal resolutions of 3.34 Å (3.79 Å against the fitted atomic model) and 4.11 Å (4.75 Å against the fitted atomic model), respectively (**Fig. S3B-E, Table S3**). The register of individual amino acids on CreS_sat_ was assigned based on the binding site of NB13 on CreS and the observed patterns of well-resolved side chain densities (**Materials and Methods, Figs. S3B**). The resulting atomic models of CreS_sat_ and NB13 allowed for interpretation of CreS_wt_ maps, determined at lower resolutions (4.36 Å −5.78 Å) (**Fig. S4, Table S3**).

**Figure 2.**
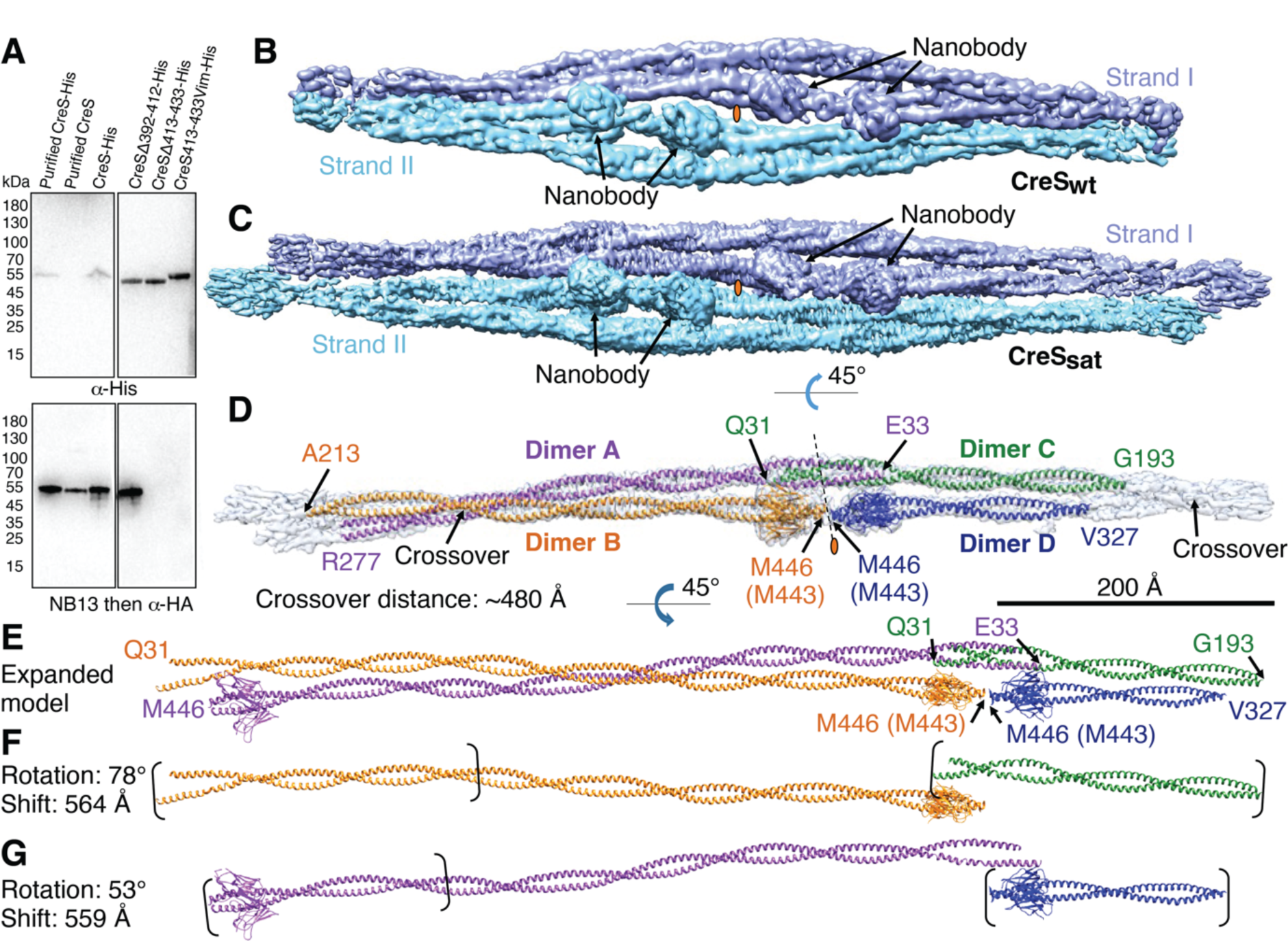
Architecture and assembly of CreS filaments. (**A**) Mapping of the CreS-specific nanobody (NB13) binding region on CreS *in vitro*. All samples except for purified proteins as indicated were cell lysates. The samples were immunoblotted with α-His for analysis of protein expression and with NB13 followed by α-HA for analysis of NB13 binding to CreS. Cryo-EM shows that CreS_wt_ (**B**) and CreS_sat_ (**C**) filaments share a common architecture formed by two strands (shown in cyan and blue) that twist around each other. (**D**) Each strand, with roughly the length of an asymmetric unit shown, consists of two CreS parallel coiled coil dimers that are held together via inter-dimer interactions. (**E**) The expanded atomic model reveals the structure of nearly complete CreS molecules, mostly missing N-terminal residues before Q31 that are presumably disordered. (**F-G**) Transformation of individual CreS dimers along the filament axis in a roughly linear fashion produces the observed single-stranded filamentous assembly. Panels **B-G** are drawn to scale (black scale bar: 200 nm). The positions of pseudo two-fold symmetry axes are indicated by an orange oval. All CreS atomic models shown are of CreS_sat_ throughout all figure panels unless otherwise stated. In these cases, residues are numbered based on CreS_sat_, whereas residue numbers in parentheses are according to CreS_wt._

### Architecture of CreS filaments

In the presence of MB13, CreS_wt_ and CreS_sat_ similarly assemble into a “supertwist”-like filament, where two intertwined strands, I and II, are related by a two-fold axis perpendicular to the filament axis (**Fig. 2B-C**). When looking at the map down this two-fold axis, each strand consists of four partial CreS dimers (dimers A, B, C, and D for strand I, and symmetry related dimers A’, B’, C’ and D’ for strand II) within a viewing window of ∼960 Å wide. Each dimer is a parallel, polar coiled coil of CreS molecules. Strands I and II are held together via interactions between CreS dimers. Taking strand I as an example, inter-dimer interactions are found between a pair of dimers near their N-termini (A-C pair) or their C-termini (B-D pair) in the longitudinal direction (i.e., along the filament axis) (**Fig. 2D**). Hence, the two-stranded CreS filament presented here lacks polarity, as is the case for the individual strands on their own. This is reminiscent of eukaryotic intermediate filaments (IFs), which are also known to be non-polar [20–23]. Furthermore, each pair of longitudinally associated dimers are structurally similar to each other, especially at the regions close to the inter-dimer interface near the N-termini or C-termini. Thus, every pair of longitudinally associated dimers are related by a local pseudo two-fold axis perpendicular to the inter-strand two-fold axis mentioned above (**Fig. 2D**). Each pair of laterally interacting dimers (A-B pair or C-D pair) alternate at a crossover due to them twisting around each other, giving rise to a crossover distance of ∼480 Å along the strand (**Fig. 2D**).

Assembling a nearly complete CreS dimer (residues 31-446 for CreS_sat_ or 31-443 for CreS_wt_), based on the observed partial dimers A and B, resulted in an expanded atomic model of the strand (**Fig. 2E**). The partial dimer C is superimposable with the equivalent part of the expanded dimer B upon a shift of 564 Å and a rotation of 78° (**Fig. 2F**). Likewise, the partial dimer D is superimposable with the equivalent region of the expanded dimer A upon a shift of 559 Å and a rotation of 53° (**Fig. 2G**). These observations indicate that every strand has a regular spacing of ∼560 Å, which corresponds to the length of a CreS molecule in a coiled coil. As a result, these transformations enable each strand and the two-stranded filament to propagate in a roughly linear fashion.

### Assembly of CreS filaments

A detailed structural analysis revealed five different types of dimer-dimer interactions that result in the observed CreS filament (**Fig. 3A**). Within each strand, there are both lateral and longitudinal interactions between CreS dimers. Types 1 and 2 involve longitudinal inter-dimer interactions near the C-termini (pairs B-D and B’-D’, residues ∼443-457, most of which are disordered in our cryo-EM maps) and near the N-termini (pairs A-C and A’-C’, residues ∼31-83), respectively (**Fig. 3B**).

**Figure 3.**
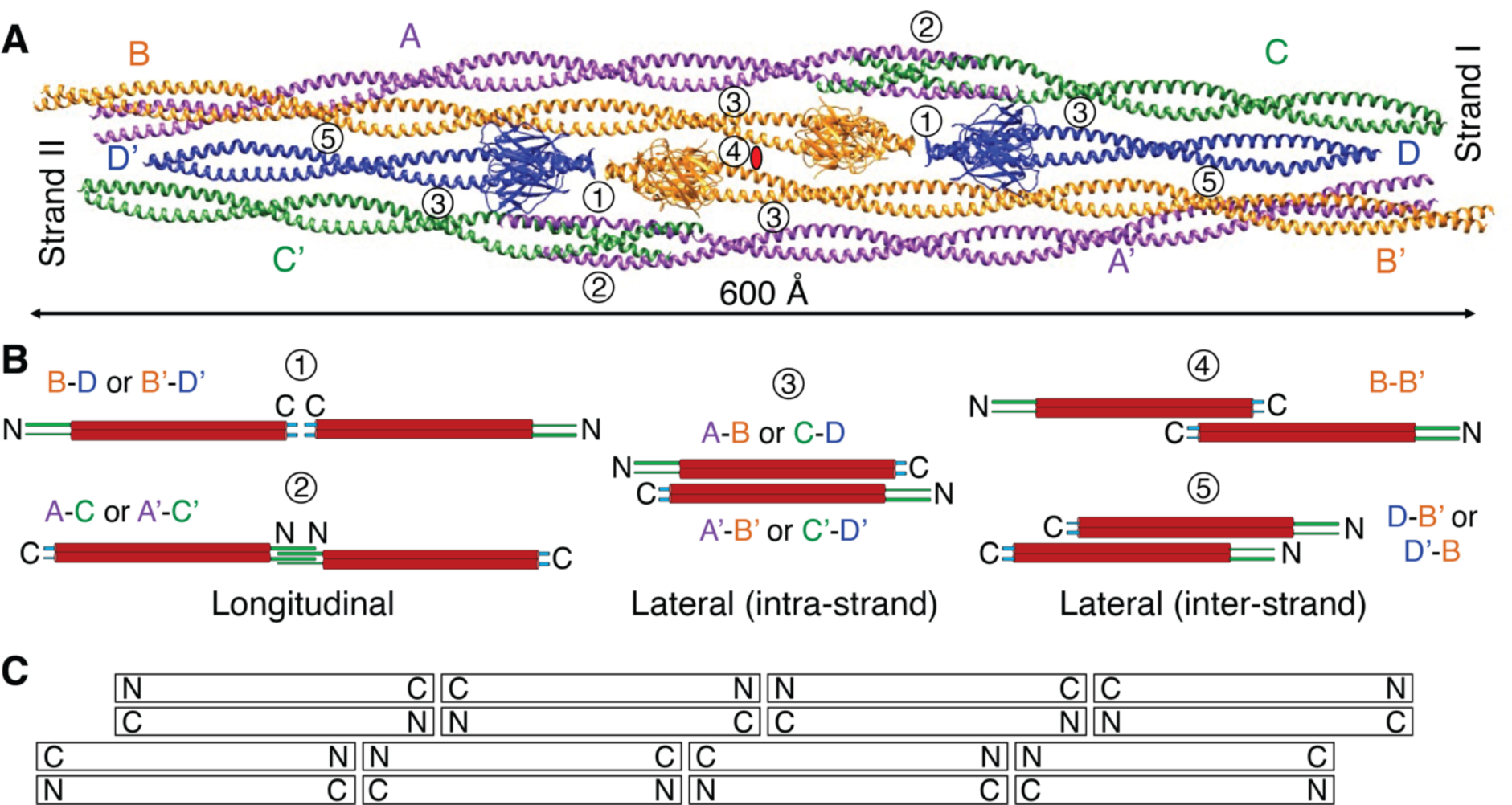
Contacts between CreS coiled coil dimers. (**A**) CreS coiled coil dimers assemble into the two-stranded filament structure via a multitude of longitudinal and lateral inter-dimer interactions that can be classified into five types. (**B**) Schematic representation of the five types of observed dimer-dimer interactions. **(C)** Schematic of the octameric CreS filament, highlighting significant differences to current intermediate filament (IF) models (see also **Fig. S5**).

Type 3 represents lateral inter-dimer interactions (pairs A-B, C-D, A’-B’, C’-D’) (**Fig. 3B**). Dimer-dimer interactions between the two strands are primarily along the lateral direction, including type 4 (pair B-B’, antiparallel) and type 5 (pairs B-D’ and B’-D, parallel) (**Fig. 3B**). Whereas relatively short stretches of amino acids are responsible for longitudinal interaction types, all lateral interaction types cover long inter-dimer binding interfaces.

As mentioned in the **Introduction**, the central rod domains of eukaryotic IF proteins are generally ∼308 amino acids long, with the exceptions of nuclear lamins and invertebrate IF proteins (which are ∼350 amino acids long). It has been proposed to be divided into three coiled coil segments with conserved lengths, 1A, 1B, and 2 [12], which are inter-connected by short linkers L1 and L12 (**Fig. S5A**). In contrast, CreS possesses a longer coiled coil rod domain (∼364 amino acids), which does not show equivalent linkers that disrupt the overall α-helical structure, based on our cryo-EM maps (**Fig. S5A**). Both N-terminal and C-terminal segments of CreS are relatively short and do not contain complex domains as seen in many eukaryotic IF proteins [24]. Furthermore, structural comparisons of CreS filamentous assemblies with those of eukaryotic IF proteins are hindered by a lack of atomic-level structural information concerning the assembly of eukaryotic IFs, despite decades of research [21, 25–28]. Nevertheless, it has been established that the coiled coil dimer of IF proteins functions as the basic subunit for IF assembly [12]. Four types of inter-dimer contacts, A11, A12, A22, and ACN, have been proposed based on information gained primarily from X-ray crystallographic studies of IF protein fragments and cross-linking mass spectrometry analyses of IF protein oligomers [29–33] [20–23] [34]. These four contact types are shown in comparison to those in CreS filaments in **Fig. S5B**, and their general lack of correspondence is elaborated on in the **Discussion**. In particular, no counterparts of the ACN type (head-to-tail) are observed in our *in vitro* CreS filament (**Fig. S5B)**. However, for eukaryotic IF proteins such as vimentin, it has been proposed that four dimers laterally assemble into an octameric protofibril, of which multiple copies come together in a symmetric way to produce the mature IF, which is often 10 nm wide [12, 26, 28, 35, 36]. The two-stranded CreS filament described here consists of four laterally associated CreS dimers that arrange in an octameric manner very loosely reminiscent of that in the proposed eukaryotic IF protofibril [26, 35] (**Fig. S5B, bottom**).

### *In vivo* crosslinking confirms crescentin filament architecture

To address the *in vivo* relevance of the CreS filament structure determined *in vitro*, we probed residue-residue contacts observed in the cryo-EM structure in *C. crescentus* cells using cysteine cross-linking with a thiol-specific and cell permeable chemical cross-linker, bismaleimidoethane (BMOE, spacer length ∼8 Å). Guided by the CreS filament structure described above, we introduced codons of pairs of cysteine residues into *creS*, which was expressed from a low-copy-number plasmid under its native promoter in *C. crescentus* cells, in a *creS* deletion background (**Materials and Methods)**. This yielded a protein level of CreS similar to the endogenous level in wt strain CB15N (**Fig. S6A**). Cells expressing wt *creS* or a *creS* mutant this way had a cell curvature comparable to wt strain, with two exceptions that will be discussed below (**Fig. S6B**). We focused on residue pairs with a Cβ-Cβ distance of 6-12 Å at multiple inter-dimer interfaces in both the CreS_sat_ and CreS_wt_ filament structures (**Table S4**).

For lateral dimer-dimer interactions (type 3, **Fig. 3**), we identified BMOE-dependent crosslinking products for at least two residue pairs in each of three regions that are well separated along the elongated inter-dimer interface (**Figs. 4 and 5**). In the absence of double-cysteine mutants, we detected no signal or only weak background signal for wt CreS and single cysteine CreS mutants, confirming that cross-linking was largely specific (**Fig. 4C**). The three lateral interaction regions probed include “crossover” (A282C/D221C, K296C/A207C, and K296C/Q204C), “middle” (T120C/R384C, A134C/K369C, and T131C/K369C), and “NC” (S34C/R419C and A37C/R419C), where NC denotes interactions between the N-terminus and C-terminus proximal regions (**Figs. 4 and 5**).

**Figure 4.**
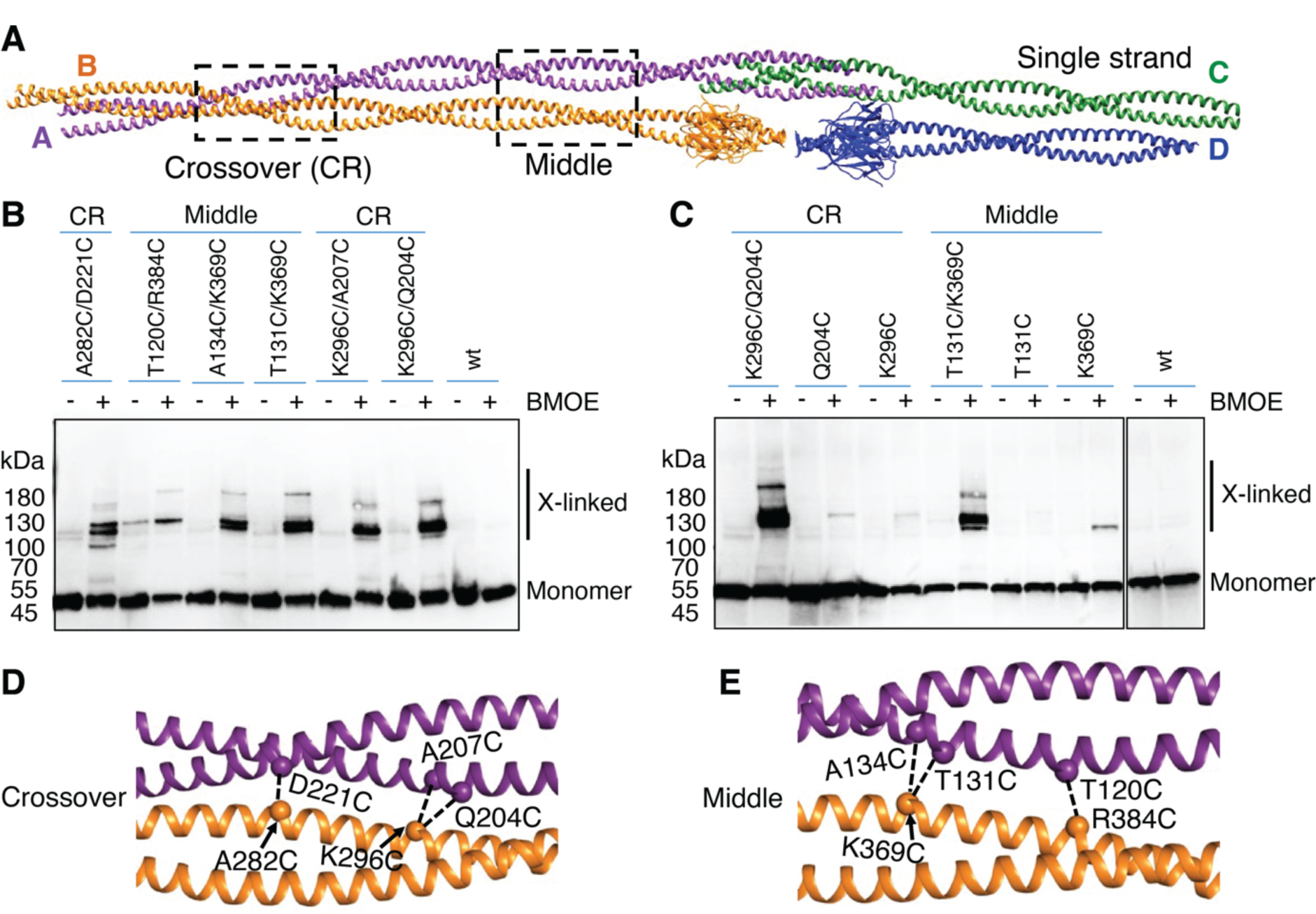
Structure-guided probing of *in vivo* CreS assembly, with a focus on lateral interactions. **(A)** The single-stranded filament structure of CreS. Dashed rectangles outline the regions in which cysteine substitutions were introduced for *in vivo* crosslinking. (**B-C**) BMOE-dependent cysteine crosslinking of Δ*creS C. crescentus* cells carrying a low-copy-number plasmid expressing *creS* or its cysteine mutants. Reaction products were subjected to Western blotting analysis with a CreS-specific nanobody, NB13. (**D-E**) A close-up view of the regions Crossover (CR, **D**) and Middle (**E**). Cα atoms are shown as spheres. Dashed lines indicate residue-residue pairs probed in **B** and **C**.

**Figure 5.**
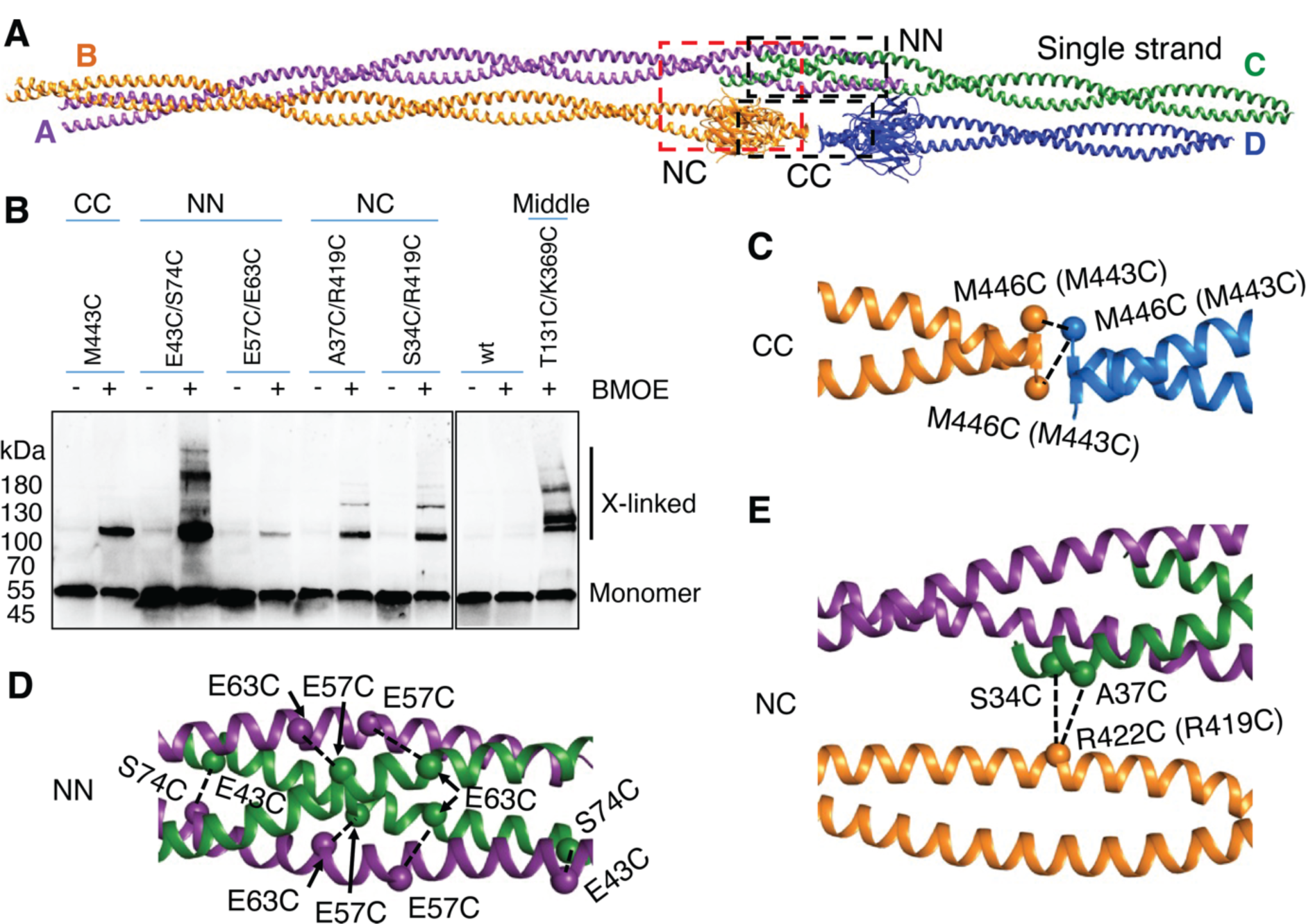
Structure-guided probing of *in vivo* CreS assembly, with a focus on longitudinal interactions. **(A)** The single-stranded structure of CreS. Dashed rectangles outline the regions where cysteine substitutions were introduced for *in vivo* crosslinking. (**B**) BMOE-mediated cysteine crosslinking of Δ*creS C. crescentus* cells carrying a low-copy-number plasmid expressing *creS* or its cysteine mutants. Reaction products were analysed as described in Fig. 4B. (**C-E**) A close-up view of the regions CC (**C**), NN (**D**), and NC (**E**). Cα atoms are shown as spheres. Dashed lines indicate residue-residue pairs probed in **B**. As in Fig. 1, residues in **C** and **E** are numbered based on CreS_sat_, whereas residue numbers in parentheses are according to CreS_wt_.

For longitudinal interactions, we observed BMOE-dependent crosslinks between residue pairs near the N-termini (E43C/S74C and E57C/E63C, region NN, type 2) and C-termini (M443C, region CC, type 1), respectively (**Fig. 5**). Since all these inter-dimer interaction regions are most likely important for the assembly of individual strands, these results indicate that the single-stranded architecture observed *in vitro* likely acts as a structural unit for producing CreS assemblies *in vivo*.

### Subcellular organization of CreS filaments

The ultrastructure of endogenous CreS assemblies at normal levels in *C. crescentus* has previously been challenging to visualize [5-8, 37, 38]. We therefore imaged wt *C. crescentus* cells where wt *creS* was moderately overexpressed using electron cryotomography (cryo-ET). Tomographic reconstructions of these cells revealed an ∼4 nm thick and on average 30-40 nm wide (minimum ∼15 nm, maximum ∼60 nm wide) structure that lines the inner cell membrane at a distance of ∼ 5 nm and that spans a major portion of the cell’s length on the concave side of the cell (**Fig. 6A**). Consistent with this structure being CreS or CreS-containing filaments, over-production of *creS_ΔN27_*, where the first 27 amino acids were truncated, in a *creS* deletion background led to prominent bundles that detach from the inner cell membrane (**Fig. 6B**). This N-terminal region is positively charged and had previously been shown to be required for attachment of CreS to the inner cell membrane [6, 38]. Additionally, similar filamentous densities were not detected in cryo-ET reconstructions of Δ*creS* cells (**Fig. 6C**). Furthermore, CTP synthase (CtpS) is another filament-forming protein that localizes to the inner cell curvature in *C. crescentus* [39]. CtpS is known to interact with CreS and acts as a regulator of cell curvature [39]. Over-production of wt *creS* in *ΔcreS* cells where *ctpS* expression was suppressed showed a similarly extended structure lining the inner cell curvature (**Fig. 6D**). Thus, CreS can form filamentous structures along the inner cell curvature in *Caulobacter* cells. This is in agreement with the subcellular localization and low-resolution views of CreS as revealed by diffraction-limited fluorescence microscopy [5–7].

**Figure 6.**
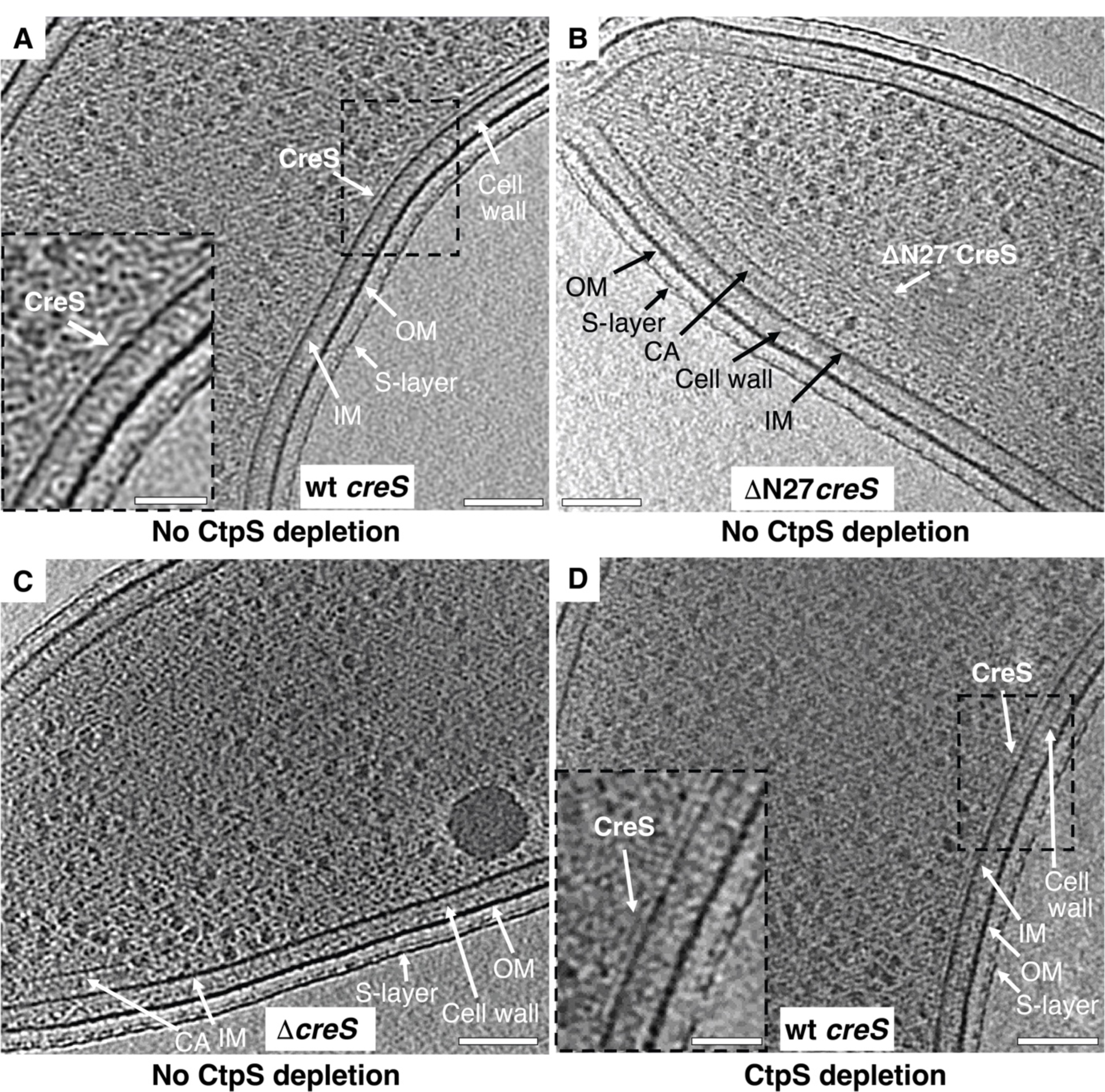
Subcellular organization of CreS filaments in cells. Tomographic slices through wild-type (wt) *C. crescentus* cells with wt *creS* overexpression (**A**, strain CJW1430), Δ*creS* cells with ΔN27 *creS* overexpression (**B**, strain YL006), Δ*creS* cells (**C**, strain LS2813), and Δ*creS* cells with both CtpS depletion and wt *creS* overexpression (**D**, strain YL062). Scale bar: 100 nm. The insets (scale bar: 200 nm) in **A** and **D** show a close-up view of the area outlined by the dashed rectangle. IM: inner cell membrane; CA: chemosensory array; OM: outer membrane.

### Functional requirements

Having determined the assembly and subcellular organization of CreS, we next addressed the molecular determinants of CreS-dependent cell curvature generation. Ectopic expression of wt *creS* in a heterologous system, *E. coli* strain C41, produced curved cells that are normally straight, as evidenced by phase contrast light microscopy and cryo-EM (**Fig. S7A, E, I and L**) [6]. Consistent with a role of N-terminal and C-terminal segments in CreS filament assembly, purified CreS_rod_ that contains only the rod domain failed to assemble into long filaments *in vitro*, and overexpression of *creS_rod_* did not curve *E. coli* cells (**Fig. S7B, F, J and L**). Further supporting this idea is that two double-cysteine mutations, E43C/S74C and E57C/E63C, in the aforementioned NN region (longitudinal inter-dimer interaction type 2) caused cell straightening in *C. crescentus* (**Fig. S6B**). Moreover, previous mutational analyses showed that deleting each of four important regions for CreS assembly observed in our structure resulted in altered assembly properties as well as partial or nearly complete loss of cell curvature in *C. crescentus* [9]. These include the rod domain (inter-dimer interaction types 3, 4, 5), the NN region (type 2), residues 121-143 (intended to correspond to the L1 linker region in eukaryotic IF proteins, types 3 and 5), and the C-terminal segment (type 1).

Nevertheless, correct CreS assembly is not the only functional requirement. Similar to previous findings using *C. crescentus* [6], deleting the first 27 amino acids, which are disordered in our cryo-EM maps, or mutating all negatively charged arginine residues to alanine residues in this region (N27RtoA) yielded straight *E. coli* cells without disrupting CreS filament assembly *in vitro* (**Fig. S7C, D, G, H, K, and L**). Thus, the faithful assembly and subcellular organization of CreS on the inner membrane are both indispensable for its function in bacterial cell shape determination.

## DISCUSSION

We have determined the complete structure of crescentin and its filament (**Figs. 1-3**). To our knowledge, this has so far not been achieved for any other IF or IF-like protein [11, 27]. Coiled coil-containing filaments are often refractory to structure determination by X-ray crystallography and also cryo-EM because of their elongated nature, the smoothness of the filaments, and heterogeneity of the filaments formed. AlphaFold 2 predictions on coiled coil proteins are not as powerful as for globular proteins [18], which is also the case for crescentin (**Fig. S1C**). To circumvent these problems, we obtained a crescentin-binding nanobody that we converted into a megabody for cryo-EM structural studies. Because the megabody bound close to the C-terminus of crescentin and all N- and C-terminal ends in the crescentin filament are located in close proximity, this enabled us to obtain a reliable structure of that part at close to 3.3 Å resolution. Further along the filament, atomic modelling was less certain, but the maps indicated that the rod domain of crescentin most likely forms a continuous coiled coil, without flexible linkers. Because we worried about the effect of the conditions used to obtain the filaments, including the use of the megabody, we used site-directed *in vivo* cysteine crosslinking to verify the obtained structure (**Figs. 4-5**). Given that we could not find any major discrepancies, we are confident that the obtained structure is a good representation of the structures crescentin forms in cells. Further supporting this notion is that mutations in a number of structurally important regions for CreS assembly identified here lead to an impaired or abolished cellular function of CreS, based on our and previous results (**Fig. S7**) [9].

Cryo-ET of cells overexpressing crescentin revealed filamentous structures on the concave sides of cells, close to the inner membrane (**Fig. 6**). Because of earlier misunderstandings [39], we demonstrated that the observed structures are not CTP synthase (CtpS). From these findings the question arises how the slowly twisting *in vitro* filaments correspond to the band-like appearance of crescentin in cells. While it is difficult to answer this without further and higher-resolution insights into the structure of the *in vivo* crescentin assemblies, it is important to note that it has been reported that when crescentin filamentous structures are released from the membrane through the inhibition of MreB or cell wall synthesis, the filaments form helical structures [6]. It seems plausible that membrane attachment removes some of the twisting that our *in vitro* structure revealed. The cryo-ET data shows crescentin to be close to the membrane (∼5 nm in distance). We propose that this distance could slow down or even stop elongasomes moving past as they circle the cells along their short axes to make cell wall [40–42].

We confirmed earlier results that the first 27 residues of crescentin are required for its membrane localization [6] (**Fig. 6**). Despite trying, we have been unable to demonstrate convincingly direct membrane binding of crescentin filaments to lipid membranes or liposomes *in vitro*. Given the above mentioned finding that the inhibition of cell wall synthesis or MreB removes crescentin from the membrane [6], it seems likely that membrane attachment, involving the first 27 residues of crescentin, is mediated by another cellular component. One such component could be MreB [7], although AlphaFold 2 is not able to predict a complex between crescentin and MreB in our hands. Possibly also speaking against this idea is the finding that crescentin over-expression in *E. coli* leads to curved cells [6]. If another component is needed to localize crescentin to the membrane, then it must be present (and functional) in *E. coli* as well.

Our structure of crescentin and its filament enables us to compare it to what is known about eukaryotic IF proteins and their filaments. It is important to point out again that there is currently no complete atomic structure of any IF protein and in particular of an IF filament [11, 26]. Given the formidable technical difficulties with these proteins, considerable efforts have been spent using a “divide and conquer” approach combining X-ray crystallography and crosslinking, to obtain pseudo-atomic models of IF proteins and filaments. Comparing our structure with these models reveals several important similarities and differences (**Fig. 3, Fig. S5**). Crescentin and most IFs form long parallel coiled coil dimers with disordered domains at each end. Vimentin and other cytoplasmic IFs have been shown to form tetramers, two dimers coming together via their coil1 domains (A11 contact) [20–23]. No such contact exists in the structure of the crescentin filament. In fact, apart from IF’s A12 contact there does not seem to be much similarity in the way dimers pack into larger fibres between IFs and crescentin. Furthermore, the most important difference between them is the longitudinal contacts. In crescentin, they are exclusively N-N (head-to-head) and C-C (tail-to-tail), whereas all the current IF models assume ACN contacts that link dimers head-to-tail. It is, however, important to point out that the experimental evidence for ACN contacts is sparse. Their existence seems to be derived from nuclear lamins that form chains of head-to-tail dimers instead of tetramers [34] and from the fact that it is difficult to build larger IF filament models from A11 tetramers without ACN head-to-tail arrangements.

All IF proteins contain a stutter that inserts into the coiled coil-forming heptad repeat four extra residues. This is thought to not interrupt the coiled coil *per se*, but to change the superhelical arrangement such that the two helices run more parallel for a short distance. Crescentin also contains a stutter (residues 406-409) and we have investigated the consequences of removing it by reverting the repeat pattern back to regular heptad repeats by inserting three residues, Ser, Ala and Thr (SAT), before position 406 [9]. While it has been reported that such a modification abolishes crescentin’s ability to curve cells [9], we have found little evidence that it changes its behaviour *in vitro*. The structures of the WT and SAT versions are very similar, except that the SAT version is slightly longer because the helices contain one almost complete extra helical turn. It will need further work to investigate this in more detail. For example, the stutter might be needed to form higher-order assemblies such as the ones seen in cells (**Fig. 6**).

Going forward it will be important to figure out if crescentin filaments locally modify and reduce cell wall synthesis via the elongasome as proposed and how crescentin filaments localize to the inner membrane since we could not reconstitute membrane binding *in vitro*. Because crescentin filaments are thin and smooth, cellular tomography is unlikely to deliver directly how this is done. Genetic and *in vitro* approaches will have to be pushed further to provide answers.

There are a number of other IF-like proteins known in bacteria, such as Scy and Filp in Streptomyces [43]. Beyond these, other bacterial coiled coil rich proteins (CCRPs), whose evolutionary relationships to IFs are even more enigmatic, such as CrvA and CrvB in *V. cholerae* and CcfM in *M. gryphiswaldense*, share a similar cellular function with CreS in maintaining curved bacterial cell shapes [44, 45]. It will be important to investigate what structures these proteins form. Revealing the evolutionary relationships between IF and their bacterial counterparts will also require more certainty about the structures of IF filaments, an area of eukaryotic structural cell biology that needs urgent attention, perhaps using similar nanobody and cryo-EM approaches as pioneered here. But even with the dissimilarities unearthed, between the crescentin filament structure and the current model of IF filaments, it is intriguing that non-polar octameric filaments made out of long parallel coiled coil dimers characterize both sets of proteins. A picture emerges in which long, extended parallel coiled coil proteins have evolved to produce a variety of filament architectures to generate mechanical and polymerization properties that are required for a multitude of cellular functions, across the tree of life.

## MATERIALS AND METHODS

### Caulobacter crescentus and Escherichia coli strains

*C. crescentus* cells were grown in peptone yeast extract (PYE) media at 30°C [46]. Antibiotics were used when required in liquid (solid) media at the following concentrations: 1 (1) µg/mL and 1 (2) µg/mL for chloramphenicol and oxytetracycline, respectively. For ectopic expression of wildtype (wt) *creS* and its variants in *C. crescentus*, a parental strain as indicated was transformed with an appropriate plasmid, based on either the low-copy-number pCT133 or medium-copy-number pCT155 vector (**Table S1**). To generate these plasmids, the sequences of *creS* and their native promoter were amplified from strain CB15N and then assembled into the pCT133 or pCT155 backbone using Gibson assembly (New England Biolabs). For protein expression in *E. coli*, an appropriate expression plasmid (**Table S1**) was electroporated into C41 (DE3) cells unless stated otherwise. Ampicillin and kanamycin were used at 100 µg/mL and 50 µg/mL, respectively, when required. *E. coli* growth and induction were both at 37°C unless stated otherwise.

For imaging, *C. crescentus* strains were grown overnight in PYE medium with the appropriate antibiotics, sub-cultured, and grown to mid-log phase. Glucose was used at a concentration of 0.2 % (w/v) to suppress protein expression driven by the leaky, xylose-inducible promoter. *E. coli* strains over-expressing CreS constructs were grown overnight in Luria-Bertani (LB) media with ampicillin to stationary phase, sub-cultured, grown to OD_600 nm_ ∼0.2, and induced by 0.05 mM IPTG for ∼2.5 h.

### Crescentin expression and production

The amino acid sequences of the proteins used in this study are listed in **Table S2**. Non-tagged wt crescentin (CreS_wt_, GenBank accession identifier: ACL97278.1) was produced in *E. coli* BL21 (DE3) cells carrying a pET29a-based plasmid for CreS_wt_ expression (strain CJW1659, C. Jacobs-Wagner, Stanford University, USA) [9]. To produce a “stutter mutant”, CreS_sat_, that has a three amino acid insertion (Ser, Ala and Thr, SAT) before residue 406 to remove the stutter, strain CJW2045 (C. Jacobs-Wagner) was used [9]. Other non-tagged CreS constructs included CreS_rod_ (amino acids 80-444), CreS_ΔN27_ (amino acids 28-457), and CreS_N274RtoA_ (all Arg mutated to Ala in amino acids 1-27). These were constructed using Gibson assembly (New England Biolabs) and employing pFE127 as a parental plasmid, which is a pHis17-based plasmid for expressing CreS-His, full length CreS followed by GSHHHHHH. The expression and purification of all non-tagged CreS proteins followed the same procedures below. All purification steps were carried out at 4°C. *E. coli* cells were grown in 2×TY media with kanamycin or ampicillin to reach an OD_600 nm_ of ∼0.6 before 1 mM IPTG was added to induce protein expression for 4 h. Cells were then spun down, and pellets were lysed by sonication in 50 mM sodium phosphate buffer, 150 mM NaCl, pH 7.2, 0.1% (v/v) Triton X-100, supplemented with lysozyme, DNase, RNase, and protease inhibitor cocktail (Roche). After centrifugation at 45,000 ×g for 40 min, the supernatant was incubated with 30 g/L of Avicel PH-10 cellulose microspheres (Sigma-Aldrich) overnight [47]. The microspheres were washed with 50 mM sodium phosphate buffer, 1 M NaCl, pH 7.2, followed by elution with 50 mM sodium phosphate buffer pH 7.2, 150 mM NaCl, 40% (w/v) glucose. The eluate was dialyzed against 50 mM CHES, pH 10, and concentrated before being injected onto a HiLoad 16/600 Superdex 200 PG column (Cytiva) for gel filtration in 5 mM Tris-HCl, pH 8. Peak fractions corresponding to CreS were pooled and concentrated using an Amicron Ultra-15 centrifugal filter (30 kDa molecular weight cutoff [MWCO], Millipore) to a final concentration of ∼9 g/L.

To purify CreS-His, cells were lysed by sonication in 50 mM Tris-HCl, 300 mM NaCl, 2 mM tris(2-carboxyethyl)phosphine (TCEP), pH 8 (buffer A), supplemented with lysozyme, DNase, RNase, and protease inhibitor cocktail (Roche). The lysate was cleared by centrifugation at 75,000 ×g for 1 h and filtered before being loaded onto a 5 mL HisTrap HP column (Cytiva). After washing with increased concentrations (0, 20, 50, and 80 mM) of imidazole in buffer A in a stepwise manner, proteins were eluted with 500 mM imidazole in buffer A. Upon concentration, the eluate was further purified via size exclusion chromatography as above, and protein fractions were treated in the same way as mentioned above.

### Expression and production of NB13 and MB13

Camelid nanobodies (NBs) were raised and selected against purified CreS_wt_ (in 5 mM Tris-HCl, pH 8) through the commercial service provided by the VIB Nanobody Core, Vrije Universiteit Brussel, Belgium. Twenty five of 97 NB clones delivered were chosen based on their sequence diversity and antigen binding capacity for another round of selection against CreS filaments at both pH 7 and pH 6.5, using an enzyme-linked immunosorbent assay (ELISA). To identify suitable NB clones for biochemical and structural studies, purified NB and megabodies (MB, a large nanobody derivative described below [13]) were produced for each of the top eight clones selected based on the ELISA data. The binding properties of each protein to purified CreS were examined using size exclusion chromatography (pH 8) and negative staining electron microscopy (pH 7 and pH 6.5). The final selection was clone No. 13, whose megabody, MB13, together with CreS, forms a stable complex at pH 8 that assembles into single, uniform filaments *in vitro* upon acidification.

To produce the nanobody version of clone No. 13, namely NB13, the non-suppressor *E. coli* strain WK6 [48] was transformed with a phagemid vector pMECS containing the PelB leader sequence, followed by the NB13 sequence and followed by an HA tag and a 6×His tag. For periplasmic expression of the megabody of NB13, MB13, with a C-terminal 6×His tag, a pET-22b-based plasmid was generated using Gibson assembly (New England Biolabs). In this construct, NB13 was grafted onto a circular permutant of the scaffold protein YgjK (*E coli* K12 glucosidase, 86 kDa) as previously described [13]. The resultant plasmid was electroporated into C41 (DE3) cells. For both NB13 and MB13, cells were grown in Terrific Broth with ampicillin, 2 mM MgCl_2_ and 0.1% (w/v) glucose until OD_600 nm_ ∼0.6. Protein expression was induced at 25°C for ∼19 h with 1 mM IPTG. All purification steps were carried out at 4°C unless stated otherwise. Cell pellets were treated with 0.2 M Tris pH 8, 0.5 mM EDTA, 0.5 M sucrose (TES) on ice for 1 h, and four times diluted TES on ice for another 1.5 h. After clearance by centrifugation at 50,000 ×g for 30 min, the resultant periplasmic extract was loaded onto a 5 mL HisTrap HP column (Cytiva). The column was washed sequentially with a) 20 mM Tris pH 8, 900 mM NaCl, b) 20 mM imidazole in buffer B (20 mM Tris-HCl, 50 mM NaCl, pH 8), c) 50 mM imidazole in buffer B, and eluted with 500 mM imidazole in buffer B. For NB13, the eluate was sufficiently pure and concentrated using an Amicron Ultra-15 centrifugal filter (3 kDa MWCO, Millipore) to a final concentration of ∼14 g/L. For MB13, the eluate was concentrated, and further purified via gel filtration using a Superdex 200 Increase 10/300 GL column (Cytiva), equilibrated in 10 mM Tris-HCl, 20 mM NaCl, pH 8. Fractions corresponding to MB13 were pooled and concentrated using a Vivaspin Turbo 15 centrifugal filter (50 kDa MWCO) to a final concentration of ∼36 g/L. Purified proteins were flash frozen in liquid nitrogen and stored at −80°C.

### Electron microscopy of negatively stained samples

Purified CreS proteins were diluted into 25 mM PIPES, pH 6.5 to reach a concentration of ∼0.2 g/L, and incubated at room temperature (RT) for 15 min to allow polymerization. The resultant sample was applied onto 400-mesh continuous carbon film copper grids (Electron Microscopy Sciences), which were then stained with 2% (w/v) uranyl acetate. Grids were imaged using a Tecnai 12 electron microscope (Thermo Fisher Scientific, TFS) equipped with an Orius CCD camera and operated at 120 kV.

### Single particle cryo-EM sample preparation

Purified CreS_wt_ or CreS_sat_ were incubated with MB13 at a molar ratio of 1:4 in 5 mM Tris-HCl pH 8 at 4°C overnight. The resultant mixture was applied onto a Superose 6 Increase 3.2/300 gel filtration column (Cytiva) in 5 mM Tris-HCl pH 8. Fractions B9 to B5, corresponding to the CreS-MB13 complex (**Fig. 1, Fig. S1**), were pooled and concentrated using a Vivaspin 500 centrifugal filter (100 kDa MWCO). The sample was diluted into 25 mM PIPES pH 6.5 to allow for polymerization at RT for 15 min, and then treated with CHAPS through the addition of a 1% (w/v) CHAPS stock solution to reach a final concentration of 0.05% (w/v) at RT for 5 min. This procedure consistently yielded uniform, single CreS filaments in complex with MB13.

For cryo-EM grid preparation, sample aliquots of 3 µL were applied onto UltrAuFoil R2/2 grids (300 mesh, Quantifoil). Grids were blotted for 1 s at 10°C with 100% relative humidity and immediately plunge frozen into liquid ethane using a Vitrobot Mark IV system (TFS). Movies of MB13-decorated CreS filaments embedded in vitreous ice were collected at liquid nitrogen temperature using a Titan Krios transmission electron microscope, operated at 300 kV and using the programme EPU (TFS). For dataset CreS_sat_, movies were collected at a nominal magnification of 81,000× in super resolution mode using a K3 direct electron detector (Gatan), resulting in a super resolution pixel size of 0.53 Å/pixel on the specimen level. The total exposure was 53 e^-^/Å^2^ in 2.4 s with a dose rate of 25 e^-^/pixel/s, a frame rate of 60 ms, and a defocus range of 0.4-2.6 µm. For dataset Cres_wt_, movies were collected at a nominal magnification of 75,000× using a Falcon 4 camera (TFS), giving a physical pixel size of 1.08 Å/pixel on the specimen level. The total exposure was 34 e^-^/Å^2^ in 10 s with a dose rate of 4 e^-^/pixel/s, a frame rate of 4 ms (EER - electron event representation), and a defocus range of 0.4-4.0 µm. Data collection statistics have been summarized in **Table S3**.

### Image processing

The image processing procedures for datasets CreS_wt_ and CreS_sat_ were essentially the same except for slight differences in pixel size and box size (see **Table S3** for details). The following description focuses on dataset CreS_sat_. Movies were corrected for inter-frame motions using MotionCor2 [49]. Aligned frames were summed and down-sampled to produce individual micrographs (1.06 Å/pixel) that were used for estimation of contrast transfer function (CTF) parameters using CTFFIND4 [50]. Nearly every single filament on the micrographs (e.g., **Fig. 1B**) showed a regular spacing of ∼60 nm between neighbouring “nodes”, of which each represents a segment of the CreS filament decorated with MB13 molecules. About 870,000 particles centred on individual nodes were picked from all micrographs that had been denoised for picking purposes using a neural network model pre-trained in Topaz [51, 52].

The following steps involved the alternate use of Relion 3.1 [53] and cryoSPARC [54]. Particles were first extracted with a box size of 100 pixels and 4.24 Å pixel size (Fourier cropping throughout the processing). About 690,000 particles were retained after reference-free 2D classification where low-quality particles were identified and removed. Six *ab initio* 3D references were then generated, and the best resolved reconstruction was used as a starting reference for global particle alignments. Particles were subsequently re-extracted with a box size of 200 pixels and 2.12 Å pixel size. Heterogenous refinement in 3D with four classes resulted in one class of particles that showed partial occupancy of MB13 molecules and those were discarded. Visual inspection of the map reconstructed using the remaining ∼600,000 particles suggested the presence of a two-fold symmetry axis perpendicular to the filament axis. Thus, the poses of these particles were then refined where C2 symmetry was imposed. Next, to produce high resolution reconstructions, particles were re-centred and re-extracted with 1.325 Å pixel size and with a box size of 320 pixels, hereafter referred to as “small box” (SB). In parallel, given that a box size of 400 Å is not sufficient to reveal the structure of a complete CreS molecule, another set of particle images were extracted with 1.59 Å pixel size and a box size of 600 pixels, hereafter referred to as “large box” (LB). The following procedures were applied independently to both sets of particle images. After 3D refinement and Bayesian polishing [55], the densities of scaffold proteins (i.e., YgjK) were subtracted from original particle images. The resultant particle images, which essentially represent nanobody-bound CreS filament segments were subjected to local refinement with C2 symmetry, yielding the reconstruction SAT-SB-C2 for set SB (SAT-LB-C2 for set LB, see **Table S3**). To improve map resolutions, particles were expanded by the C2 symmetry [56], and the densities corresponding to one of two symmetry-related copies were subtracted from images. Subsequent local refinement with C1 symmetry led to a higher resolution reconstruction, SAT-SB-C1 for set SB (SAT-LB-C1 for set LB). The resolutions of reconstructions were estimated based on two methods. One was the Fourier Shell Correlation (FSC) between two independently calculated half maps (Gold Standard FSC) using an FSC cut-off of 0.143 [57]. The other was a model-map FSC between the final EM map and a map computed based on an atomic model (described below), built into and refined against the EM map, using an FSC cut-off of 0.5 [58]. All maps were sharpened using a deep learning-based programme, DeepEMhancer [59]. Local resolution was assessed using Relion [53].

### Atomic model building and refinement

The reconstructions SAT-SB-C1 and SAT-LB-C1 showed a structure consisting of four partial CreS coiled coil dimers (i.e., two pairs of longitudinally or laterally associated partial dimers) (**Fig. 2**). The two partial dimers within each longitudinal pair are related by pseudo two-fold symmetry and are held together by interactions between the C-termini or between the N-termini. Thus, each lateral pair is formed by two segments, the N- and C-segments, with their N-terminus and C-terminus oriented towards the pseudo two-fold symmetry axis, respectively. For each segment, we computationally determined the amino acid register by aligning the observed map density with the amino acid sequence of CreS_sat_ using a previously described approach [60]. In this process, we focused on a fragment of ∼20 amino acids that showed the most prominent side chain densities in SAT-SB-C1. Briefly, every amino acid position in the fragment was assigned a number among 0-6 based on the estimated size of the side chain density by visual inspection. The resultant sequence of numbers was searched against that of the entire amino acid sequence of CreS_sat_, according to a scoring matrix, where a mismatch of a small residue in the sequence with the observed large side chain density at a given position was penalized. The amino acid register determined *in silico* this way was further supported by several lines of evidence. First, the resultant amino acid sequence fitted well into the observed side chain density (e.g., **Fig. S3**), as judged by visual inspection. Second, the binding region of NB13 on CreS was mapped to residues 413-433 (**Fig. 2A**), consistent with the assigned amino acid register for the C segment. Third, whole cell cysteine crosslinking revealed residue-residue contacts that were consistent with the atomic model of the single-stranded structure (**Figs. 4-5**). In particular, laterally associated inter-dimer residue-residue contacts supported the assigned amino acid register for the N segment (**Fig. 4**).

To interpret the highest resolution reconstruction SAT-SB-C1, we predicted the coiled coil dimeric structures for the N segment and C segment based on their estimated length using CCFold [61]. A homology model of NB13 was generated using SWISS-MODEL [62] with the structure of a nanobody (NB-ALFA) against the ALFA-tag (PDB entry 6I2G) as a template. These starting atomic models were fitted into the cryo-EM map in Chimera [63], followed by manual model re-building in Coot [64]. In this process, the orientation of NB13 was initially determined based on the β-strands identified in the map and was similar to that of NB-ALFA when complexed with the ALFA-tag that forms an α-helix. The atomic coordinates were subsequently refined against the map in real space using Phenix [65, 66], where secondary structure restraints were applied. Multiple cycles of manual model rebuilding and real space refinement improved the fitting of the model into the density map and model geometry. To interpret the reconstruction SAT-LB-C1, we expanded the atomic model for SAT-SB-C1 by fitting predicted structures of the remaining regions of CreS_sat_. In local regions where the map resolution was not sufficient for an unambiguous secondary structural assignment between a helix and a loop, we assumed that it was helical according to coiled coil and secondary structure predictions. The expanded atomic model was rebuilt and refined using the same procedures described above. For each of the C2 reconstructions, SAT-SB-C2 and SAT-LB-C2, the atomic model for their C1 counterpart and the C2 symmetry related copy were fitted separately into the map as rigid bodies, and then combined into a single atomic model. Per-atom real space refinement of the resultant atomic model was not performed due to the low-resolution nature of these reconstructions compared with their respective C1 counterparts.

The omission of three-amino acids (SAT) in CreS_wt_ with respect to CreS_sat_ produces a stutter that locally disrupts the continuity of the dimeric heptad-repeat coiled coil [67]. Thus, it is reasonable to assume that the preceding and later segments along the coiled coil have little structural changes upon introduction of the stutter. In line with this, we found that the overall arrangement of CreS_wt_ dimers in the filament is similar to that observed for CreS_sat_ (**Fig. 2**). We determined the amino acid register for the C segment using NB13 molecules as a spatial reference and for the N segment by assuming that CreS_wt_ and CreS_sat_ share the same lateral, inter-dimer interactions between the N-proximal region and C-proximal region. To interpret the reconstruction WT-SB-C1, we used the CreS_sat_ structure as a starting atomic model and replaced the segment corresponding to the stutter region in CreS_wt_ with a sequence adjusted, stutter-containing fragment in vimentin (PDB entry 1GK4). The resultant atomic model was subjected to multiple rounds of model rebuilding and real space refinement against the map, as mentioned above. For the reconstruction WT-LB-C1, the atomic model for WT-SB-C1 was expanded with appropriate segments from the CreS_sat_ structure, followed by model rebuilding and refinement. The atomic model for each of the two C2 reconstructions, WT-SB-C2 and WT-LB-C2, was generated using the aforementioned rigid body fitting approach.

The final atomic models were geometrically validated based on the criteria of MolProbity [68]. Model statistics have been summarized in **Table S3**. All figures were generated using Pymol (https://pymol.org/) or Chimera [63].

### Electron cryotomography (cryo-ET)

*C. crescentus* cells were pelleted at 8,000 ×g for 2 min and resuspended in PYE media to reach an OD_600 nm_ of ∼5. The resuspension was mixed with protein A-conjugated 10 nm gold fiducials (BBI Solutions). Aliquots of 2.5 µL sample were applied onto freshly glow-discharged Quantifoil R 3.5/1 Cu/Rh (200 mesh) grids, followed by blotting for 2-3 s and plunge freezing into liquid ethane using a Vitrobot Mark IV system (TFS). Specimens were imaged using a Titan Krios transmission electron microscope (TFS), operated at 300 kV and equipped with a BioQuantum imaging filter (Gatan) and a K3 camera (Gatan). Tilt series were collected from −60° to 60° in 2° increments at a nominal magnification of 19,500× with a pixel size of 3.84 Å /pixel using SerialEM [69]. The total exposure was ∼180 e^-^/Å^2^ with a defocus of 8-10 μm. Tilt series were aligned automatically using Batchruntomo in IMOD [70, 71], and tomograms were reconstructed using the SIRT algorithm in TOMO3D [72].

### Mapping of the NB13 binding site on crescentin

To map the binding site of NB13 on CreS, CreS-His variants were constructed for ectopic expression in C41 (DE3) cells by introducing either a truncation or a substitution into the parental pFE127 plasmid via site-directed PCR-based mutagenesis (KLD enzyme mix, New England Biolabs). Protein expression was induced for 4 h at 37°C by adding 1 mM IPTG when cell cultures reached OD_600 nm_ ∼0.5 in LB with ampicillin. Cell pellets were washed twice with cold PBS and lysed using CelLytic Express (Sigma-Aldrich) with protease inhibitor cocktail (Roche) at RT for 20 min. Purified CreS controls and equivalent OD_600 nm_ units of cleared cell lysates by centrifugation were resolved on a 4-20% SDS-polyacrylamide gel (Bio-Rad). In this step, each biological sample was split into two halves, and the two halves were loaded separately on the same gel. For Western blotting analysis, one half was immunoblotted with NB13 (7 µg/mL) followed by washing and probing with α-HA-peroxidase (1:1000, Roche), whereas the other half was probed with α-His-peroxidase only (1:4000, TFS).

### Whole cell cysteine crosslinking

*C. crescentus* strains harbouring a low-copy-number plasmid for expressing CreS or its mutants were grown overnight in PYE with oxytetracycline, sub-cultured, and grown to OD_600 nm_ 0.5-0.6. Control strains, CB15N and CB15N Δ*creS* were grown in the same way without antibiotics. About 1.2 OD_600 nm_ units of cells were spun down at 8,000 ×g for 3 min, and then kept on ice for the following steps. Pellets were washed with cold PBS and resuspended in 80 µL of cold PBS, followed by a 10 min reaction upon addition of 2 µL DMSO or bis(maleimido)ethane (BMOE, final concentration 0.5 mM, ∼8 Å arm length, TFS) in DMSO. The reaction was quenched by adding 1 µL of β-mercaptoethanol (BME, stock ∼2.3 mM). Cells were then lysed using CelLytic Express (Sigma-Aldrich), supplemented with protease inhibitor cocktail (Roche) at RT for 15 min. The suspension was incubated at 70°C for 5 min in the presence of LDS loading buffer supplemented with 4% (v/v) BME. Samples equivalent to 0.1 OD_600 nm_ units of cells were resolved on a 4-20% SDS-polyacrylamide gel (Bio-Rad), and immunoblotted with purified NB13 (16 µg/mL), followed by washing and then probing with α-HA-peroxidase (1:1000, Roche).

### Light microscopy

Cells grown in conditions described above were imaged upon immobilization on 1% agarose pads. Images were acquired using a Nikon Eclipse Ti2 microscope equipped with a Neo sCMOS camera (Andor) and a Nikon Plan APO DIC objective (100×, numerical aperture 1.40). Cell segmentation and cell curvature analysis were performed using MicrobeJ [73] as a plugin in ImageJ [74]. The number of cells analysed was between 114 and 519 for *C. crescentus* strains, and between 140 and 293 for *E. coli* strains. For each strain, mean cell curvature (µm^-1^) and standard deviation (µm^-1^) were computed and presented.

## DATA AVAILABILITY

The atomic coordinates derived using reconstructions SAT-SB-C2, SAT-SB-C1, SAT-LB-C2, SAT-LB-C1, WT-SB-C2, WT-SB-C1, WT-LB-C2, and WT-LB-C1 have been deposited with the Protein Data Bank (accession numbers 8AFH, 8AFE, 8AJB, 8AHL, 8AFM, 8AFL, 8AIX, 8AIA). The cryo-EM maps of SAT-SB-C2, SAT-SB-C1, SAT-LB-C2, SAT-LB-C1, WT-SB-C2, WT-SB-C1, WT-LB-C2, and WT-LB-C1 have been deposited with the Electron Microscopy Data Bank (accession numbers EMD-15398, EMD-15395, EMD-15476, EMD-15446, EMD-15402, EMD-15401, EMD-15473, EMD-15465). Table S3 summarises details.

## AUTHOR CONTRIBUTIONS

Y.L. and J.L. designed research; Y.L performed most research, F.v.d.E made CreS constructs; Y.L. and J.L. analysed data; Y.L. and J.L. wrote the manuscript.

## ACKNOWLEDGEMENTS AND FUNDING SOURCES

We thank all members of the MRC Laboratory of Molecular Biology (Cambridge, LMB) electron microscopy facility for excellent EM support, all members of the LMB light microscopy facility for technical support, Diamond Light Source (Harwell, UK) for access to, and support at the cryo-EM facilities at the UK’s National Electron Bio-imaging Centre (eBIC), funded by the Wellcome Trust, MRC and BBSRC, and Toby Darling and Jake Grimmett (LMB Scientific Computing) for support. We are grateful to Lucy Shapiro (Stanford University, USA), Christine Jacobs-Wagner (Stanford University, USA), Zemer Gitai (Princeton University, USA), and Michael Laub (MIT, USA) for sharing *C. crescentus* and *E. coli* strains and plasmids used in this work. We thank all members of the Löwe group for helpful discussions. This work was funded by the Medical Research Council, UK (U105184326 to J.L.) and the Wellcome Trust (202754/Z/16/Z to J.L.).

## SUPPLEMENTARY FIGURES

**Figure S1.**
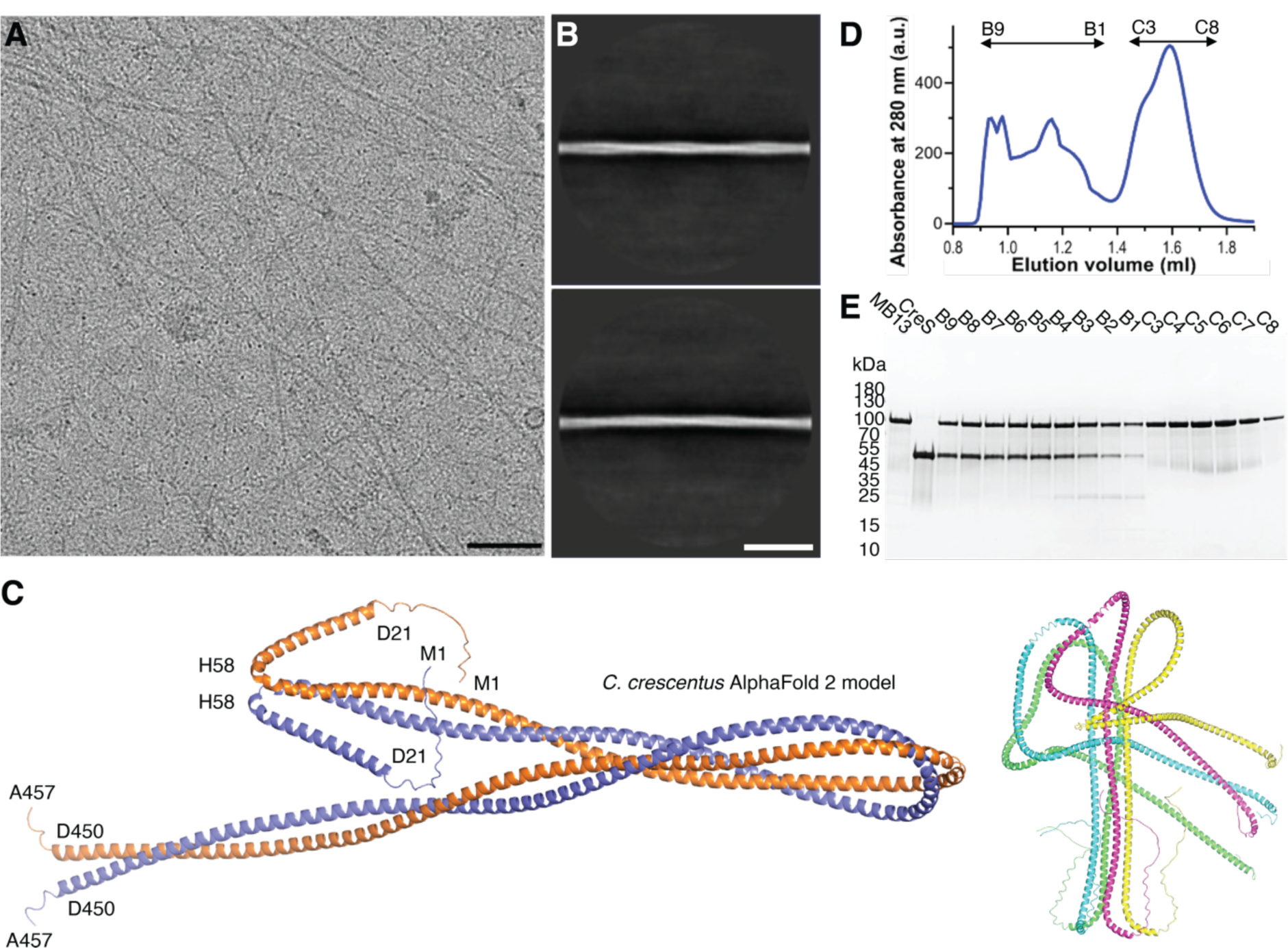
CreS_wt_ assembles into two-stranded filaments *in vitro*. (**A**) Cryo-EM micrograph of single filaments of CreS_wt_ formed at pH 7. (**B**) Two-dimensional (2D) class averages show the presence of two intertwined strands. Scale bar: 100 nm (**A**) and 50 nm (**B**). Megabody MB13 forms a complex with CreS_wt_. **(C)** Left: annotated AlphaFold 2 model of CreS dimer. Right: AlphaFold 2 model of CreS tetramer. (**D**) Size exclusion chromatography (SEC) profile of CreS_wt_, preincubated with an excess of MB13 (1:4 molar ratio) at pH 8. (**E**) SDS-PAGE analysis of SEC fractions. Fractions B9 to B5 were pooled and used for EM analysis. MB13 top, CreS_wt_ bottom bands.

**Figure S2.**
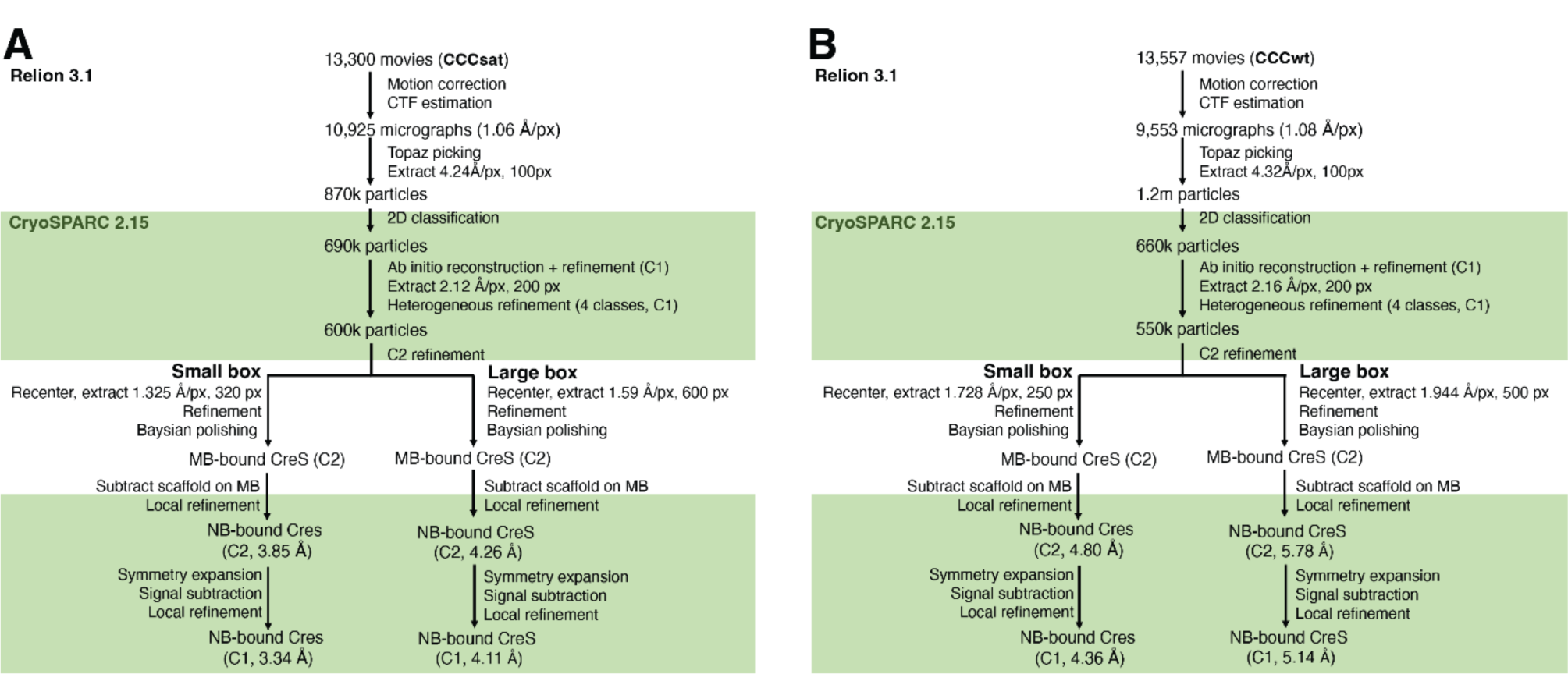
Cryo-EM structure determination of CreS. Image processing workflow used for CreS_sat_ (**A**) and CreS_wt_ (**B**).

**Figure S3.**
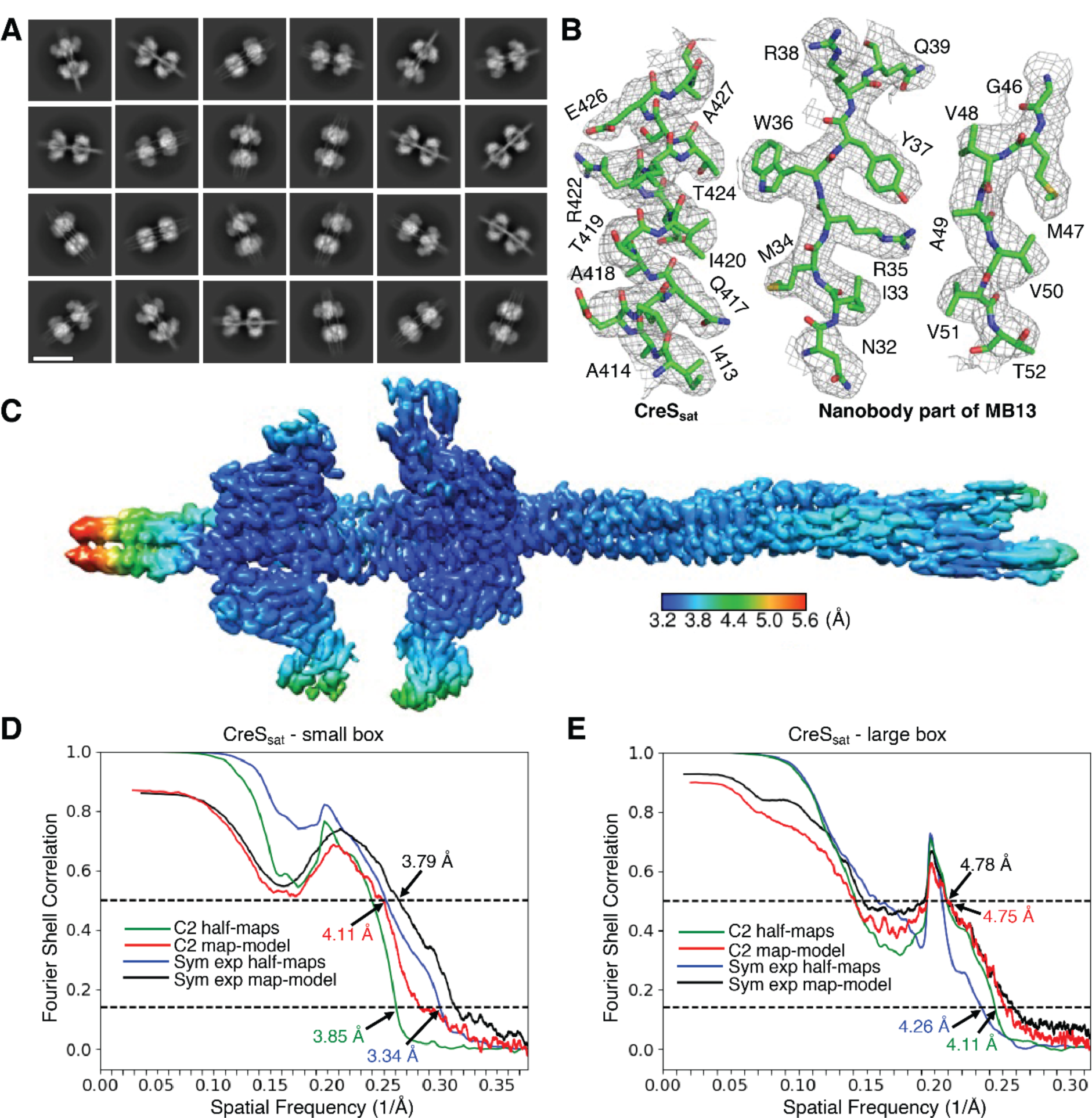
Cryo-EM analysis of CreS_sat_ filaments in complex with megabody MB13. Related to Figure 2. (**A**) 2D class averages of individual nodes formed by MB13 and CreS_sat_ in different orientations. (**B**) Typical cryo-EM map densities for CreS_sat_ and MB13, with the fitted atomic model superimposed. (**C**) Local resolution of the symmetry-expanded map reconstructed with a box size of 424 Å. FSC-based resolution estimation for maps reconstructed using a box size of 424 Å (**D**, small box) and 960 Å (**E**, large box).

**Figure S4.**
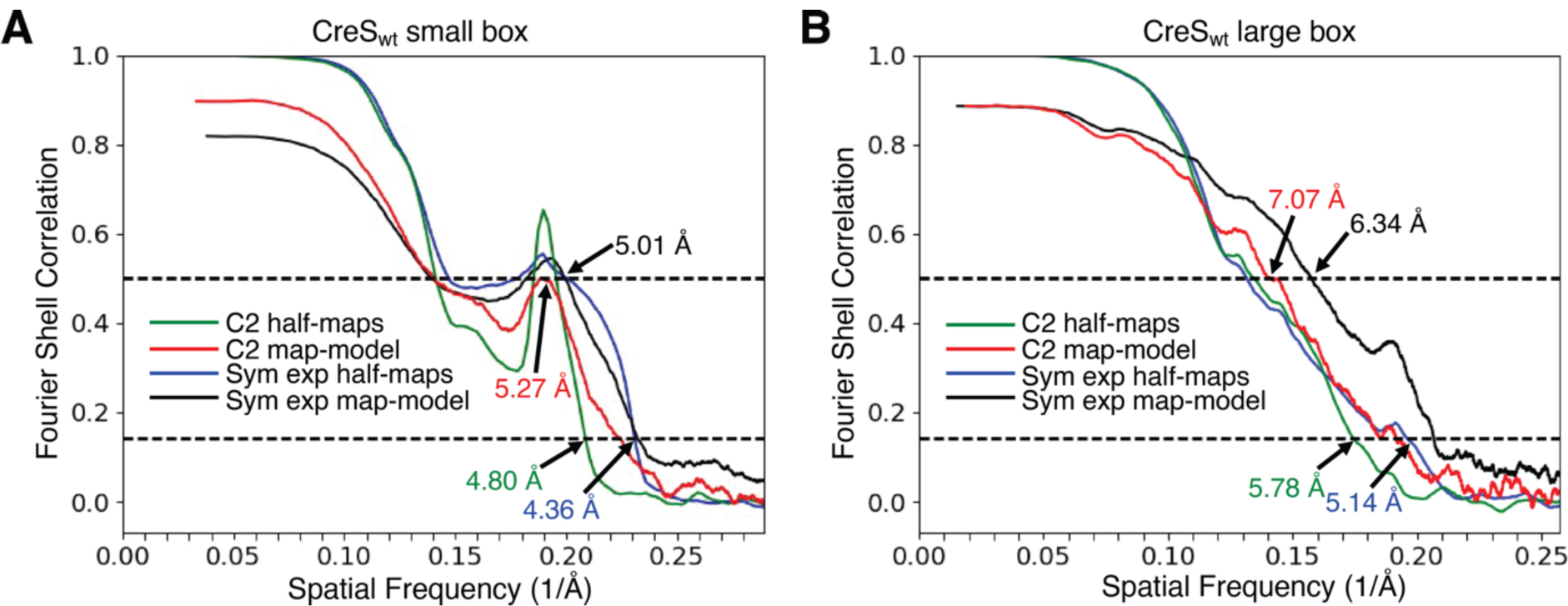
Cryo-EM analysis of CreS_wt_ filaments in complex with megabody MB13. Related to Fig. 2. Resolution estimation for maps reconstructed using a box size of 432 Å (**A**, small box) and 972 Å (**B**, large box) based on FSC curves.

**Figure S5.**
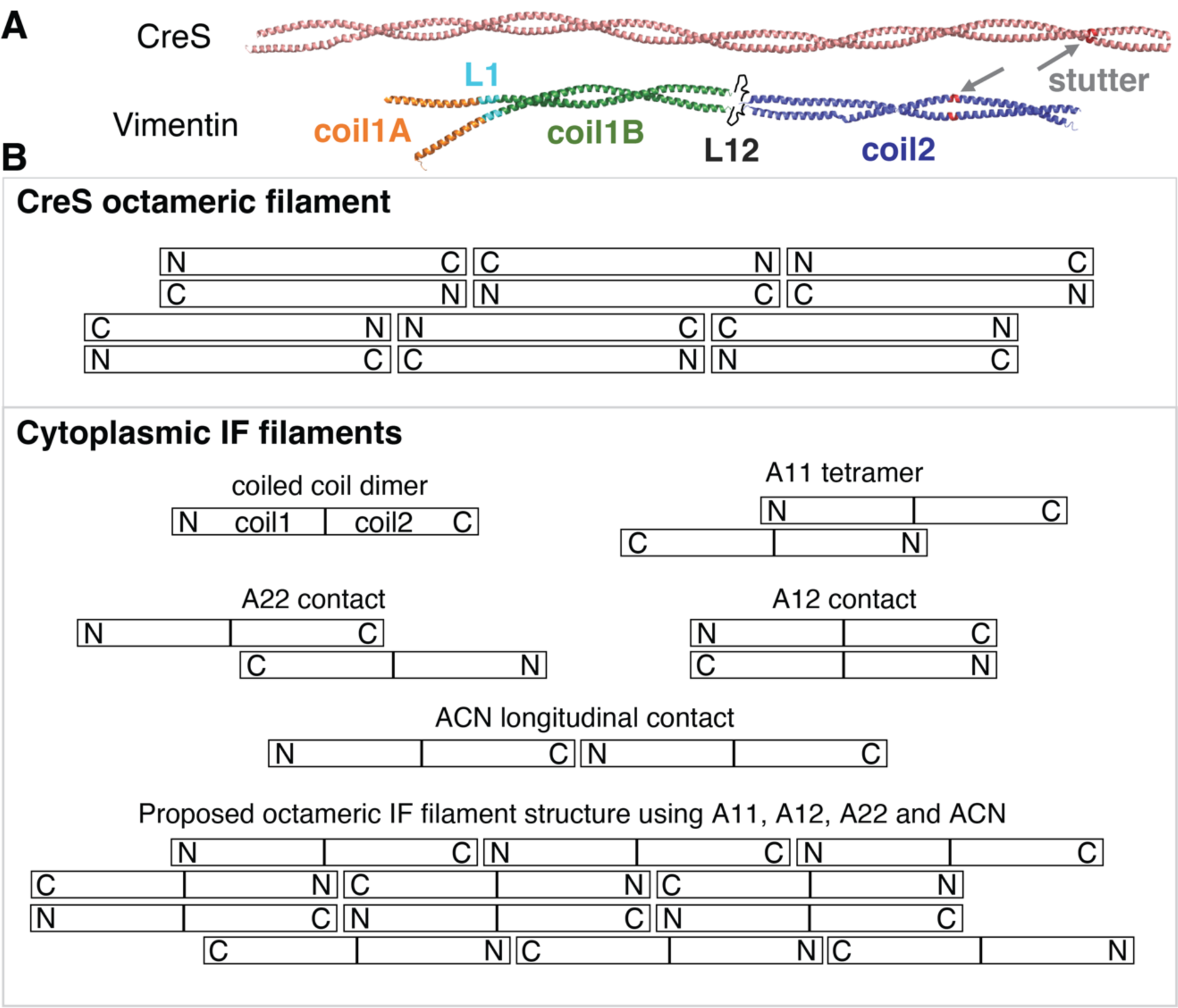
Structural comparison of CreS with eukaryotic IF proteins. (**A**) To-scale comparison of the atomic model of a CreS dimer (upper) with the composite atomic model of a vimentin dimer (lower). The IF structure shown is a hybrid model assembled from crystal structures of vimentin fragments (PDB entries 1GK4, 3S4R, 3TRT, 3UF1, and 1GK6), similar to what was reported [21]. (**B**) Comparison of crescentin filaments (upper box, same as Fig. 3C) with current models of cytoplasmic IF protein filaments (lower box). IF proteins also form parallel and extended coiled coil dimers. Many IF proteins have been shown to form staggered and antiparallel “A11” tetramers, in which two dimers come together via their coil1 rod segments. IF polymers have also been shown, mostly by cross-linking studies, to contain staggered A22 and flush A12 contacts. End-to-end contacts are believed to be via N- and C-terminal ends coming together, head-to-tail ACN contacts. There is little experimental evidence for ACN contacts in cytoplasmic IF filaments, but it is difficult to build gap-free models using the other contacts, without ACN contacts. The current model of an IF “octameric” protofilament is shown at the bottom [11]. The most striking difference to the structure of crescentin filaments discovered here is that all longitudinal contacts are head-to-tail (N-C) in IFs (ACN), and not N-N and C-C head-to-head and head-to-tail, as in crescentin. Both filaments lack overall polarity and one inter-dimer contact is similar, A12 and interaction type 3 **(Fig. 3A & C)**, whereas the others are different. For example, there is no parallel inter-dimer contact in the IF model, as in crescentin’s interaction type 5 **(Fig. 3A & C)**. Furthermore, A11 and A22 in IF filaments are more substantial than the somewhat related contact types 2 (N-N) and 4 (C-C overlap) in crescentin, respectively **(Fig. 3A & C)**. While somewhat related in overall architecture, the way the dimers come together to form filaments can be described as significantly different.

**Figure S6.**
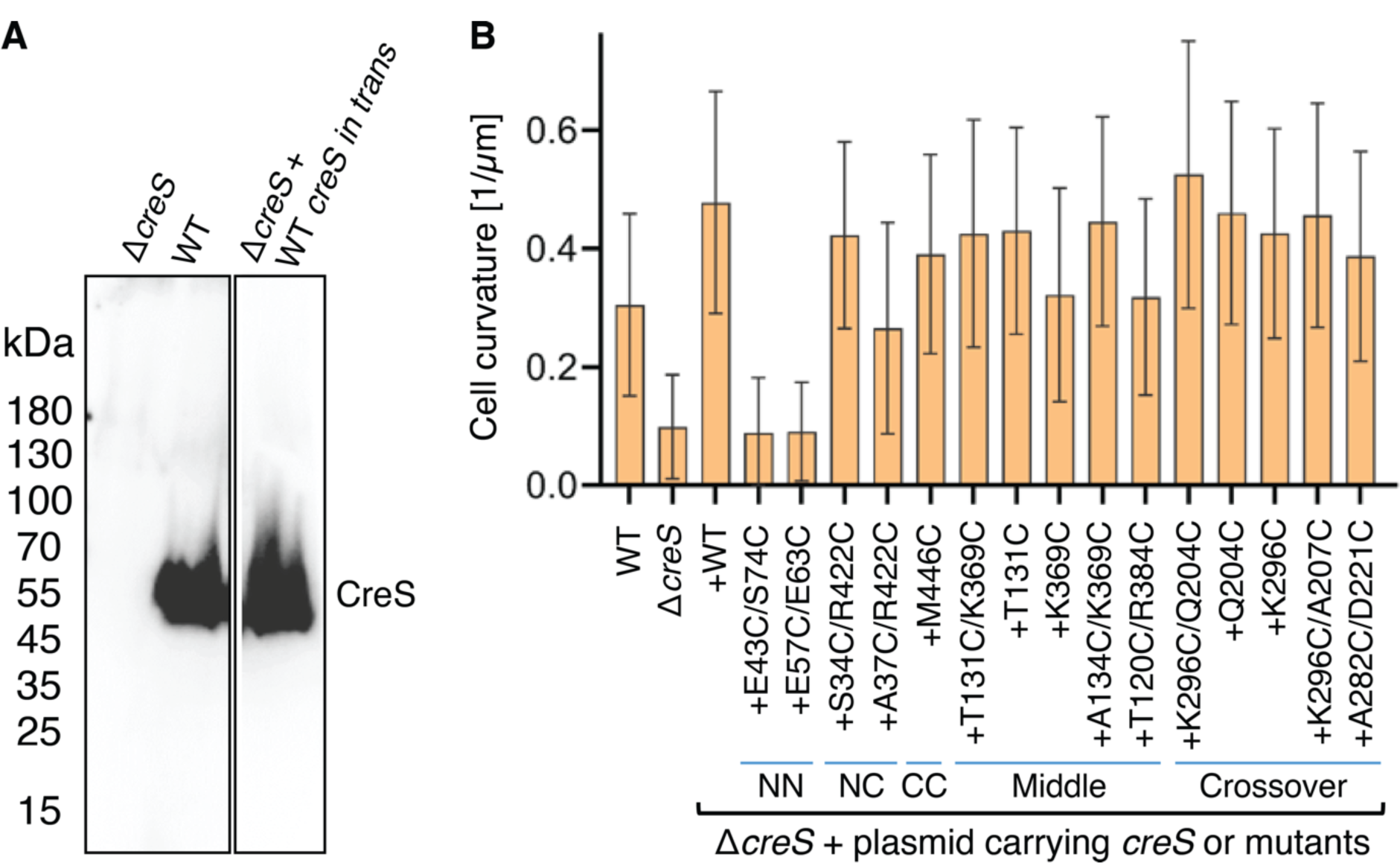
Characterization of *C. crescentus* cells carrying a *creS*-expressing plasmid. (**A**) Western blotting analysis of CreS protein levels in *C. crescentus* strains. Production of CreS from a low-copy-number plasmid under its native promoter in a *creS* deletion background yields a similar protein level as that in the wt strain. (**B**) Quantification of cell curvature of *C. crescentus* strains used for cysteine crosslinking experiments (as in **Figs 4-5**). The mean curvature values are shown, and error bars represent standard deviations. The number of cells analysed for each strain was between 114 and 591.

**Figure S7.**
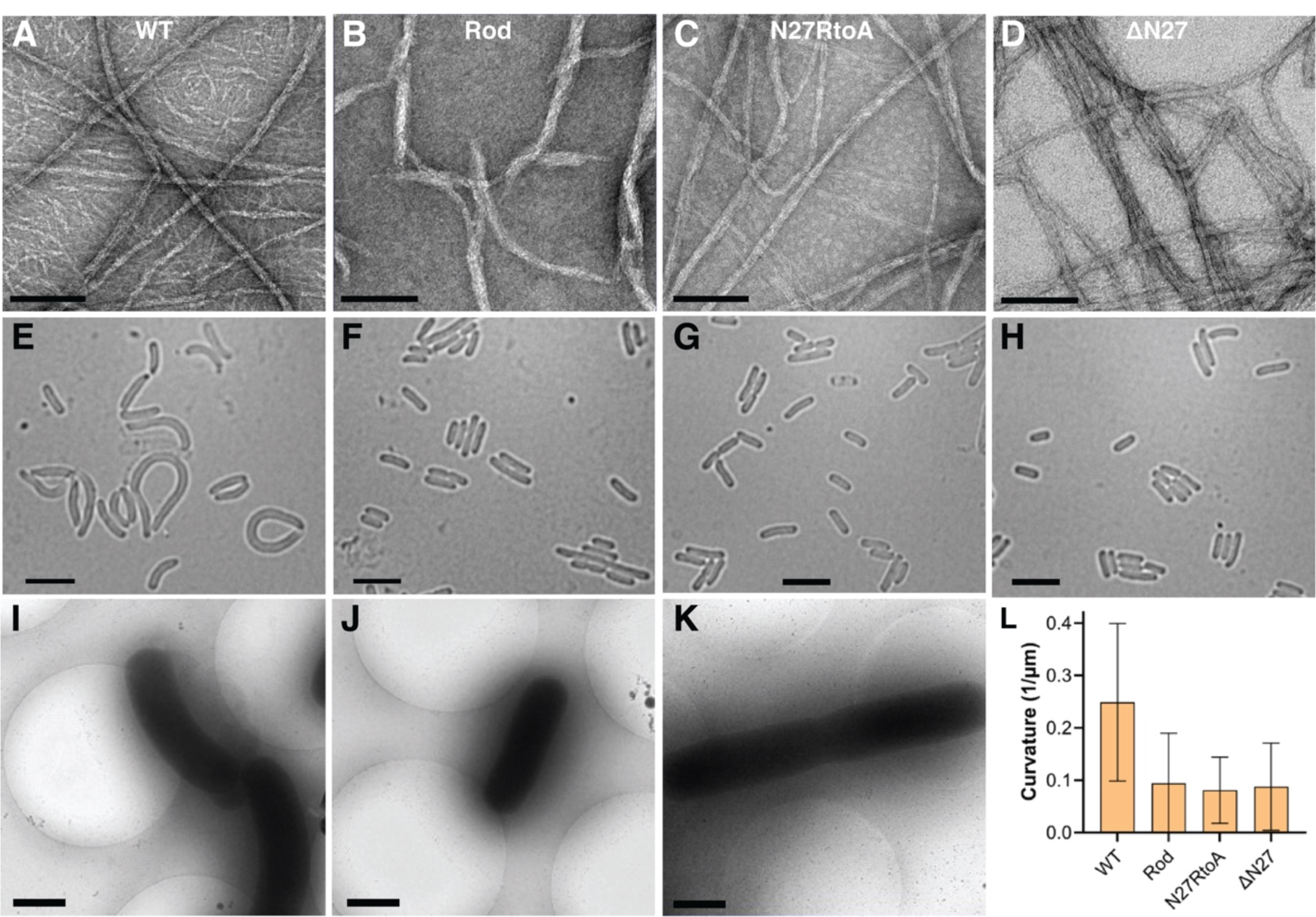
The N-terminal residues 1-27 of CreS are required for its cell shape function in *E. coli*. (**A-D**) EM micrographs of negatively stained filaments formed *in vitro* using purified CreS_wt_ (**A**), CreS_rod_ (**B**), CreS_N27RtoA_, where all Arg residues were substituted with an Ala in the N-terminal 27 amino acids (**C**), and CreS_ΔN27_ (**D**). Scale bar: 100 nm. (**E-J**) Typical phase contrast images of *E. coli* cells overexpressing CreS_wt_ (**E**), CreS_rod_ (**F**), CreS_N27RtoA_ (**G**), and CreS_ΔN27_ (**H**). Scale bar: 5 µm. (**I-K**) Typical cryo-EM images of *E. coli* cells overexpressing CreS_wt_ (**I**), CreS_rod_ (**J**), and CreS_N27RtoA_ (**K**). Scale bar: 1 µm. (**L**) Quantification of cell curvature based on phase contrast images (corresponding to panels **E-J**). Shown are the mean curvature values with error bars representing standard deviations. The numbers of cells analysed for each strain were between 140 and 293. Protein expression was induced by 0.05 mM IPTG for ∼2.5 h.

## SUPPLEMENTARY TABLES

**Table S1.** Strains and plasmids.

| Name | Description/Genotype | Source |
| --- | --- | --- |
| **Strains** |  |  |
| *E. coli* |  |  |
| BL21 (DE3) | F<sup>-</sup> *ompT hsdSB (rB- mB-) gal dcm* (DE3) | NEB |
| C41 (DE3) | F<sup>-</sup> *ompT hsdSB (rB- mB-) gal dcm* (DE3) | Sigma-Aldrich |
| WK6 | F' *lacIq Δ(lacZ)M15 proA+B+ Δ(lac-proAB) galE rpsL* | VIB |
| CJW1659 | BL21 (DE3) expressing CreS<sub>wt</sub> (non-tagged) | [1] |
| CJW1661 | C41 (DE3) expressing CreS<sub>wt</sub> (non-tagged) | [1] |
| CJW2045 | BL21 (DE3) expressing CreS<sub>sat</sub> (insertion of three amino acids (Ser, Ala, SAT, at position 406) | [1] |
| *C. crescentus (vibrioides)* |  |  |
| CB15N | Synchronizable strain of CB15 | [2] |
| LS2813 | CB15N Δ*creS* | [3] |
| ZG216 | CB15N Δ*creS* *ctpS*<sup>1-400</sup> P<sub>xyI</sub>-*ctpS* (*ctpS* truncated to first 400 bp and another *ctpS* under a xylose inducible promoter on chromosome) | [4] |
| CJW1430 | CB15N + pJS14-based medium-copy-number plasmid for expressing CreS under its native promoter | [5] |
| YL010 | CB15N Δ*creS* + pCT133-P<sub>creS</sub>-CreS<sub>wt</sub> | This study |
| YL006 | CB15N Δ*creS* + pCT133-P<sub>creS</sub>-CreS<sub>ΔN27</sub> | This study |
| YL012 | CB15N Δ*creS* + pCT133-P<sub>creS</sub>-CreS<sub>M443C</sub> | This study |
| YL017 | CB15N Δ*creS* + pCT133-P<sub>creS</sub>-CreS<sub>E43C+S74C</sub> | This study |
| YL018 | CB15N Δ*creS* + pCT133-P<sub>creS</sub>-CreS<sub>E57C+E63C</sub> | This study |
| YL019 | CB15N Δ*creS* + pCT133-P<sub>creS</sub>-CreS<sub>R419C+S34C</sub> | This study |
| YL020 | CB15N Δ*creS* + pCT133-P<sub>creS</sub>-CreS<sub>R419C+A37C</sub> | This study |
| YL022 | CB15N Δ*creS* + pCT133-P<sub>creS</sub>-CreS<sub>K369C+T131C</sub> | This study |
| YL023 | CB15N Δ*creS* + pCT133-P<sub>creS</sub>-CreS<sub>K369C+A134C</sub> | This study |
| YL024 | CB15N Δ*creS* + pCT133-P<sub>creS</sub>-CreS<sub>R384C+T120C</sub> | This study |
| YL025 | CB15N Δ*creS* + pCT133-P<sub>creS</sub>-CreS<sub>K296C+Q204C</sub> | This study |
| YL026 | CB15N Δ*creS* + pCT133-P<sub>creS</sub>-CreS<sub>K296C+A207C</sub> | This study |
| YL027 | CB15N Δ*creS* + pCT133-P<sub>creS</sub>-CreS<sub>A282C+D221C</sub> | This study |
| YL031 | CB15N Δ*creS* + pCT133-P<sub>creS</sub>-CreS<sub>Q204C</sub> | This study |
| YL032 | CB15N Δ*creS* + pCT133-P<sub>creS</sub>-CreS<sub>K296C</sub> | This study |
| YL033 | CB15N Δ*creS* + pCT133-P<sub>creS</sub>-CreS<sub>T131C</sub> | This study |
| YL034 | CB15N Δ*creS* + pCT133-P<sub>creS</sub>-CreS<sub>K369C</sub> | This study |
| YL062 | ZG216 + pCT155-P<sub>creS</sub>-CreS<sub>WT</sub> | This study |
| **Plasmids** |  |  |
| pCT155-P<sub>van-3xM2-didA</sub> | pCT155-based parental plasmid for cloning (medium copy number, pJS14-based, chlor<sup>R</sup>) | [6] |
| pCT133-P<sub>van-3xM2-didA</sub> | pCT133-based parental plasmid for cloning (low copy number, pMR20-based, tet<sup>R</sup>) | [6] |
| pHis17-CreS<sub>ΔN27</sub> | pHis17-based plasmid for expressing CreS<sub>ΔN27</sub> (residues 28-457) | This study |
| pHis17-CreS<sub>N27RtoA</sub> | pHis17-based plasmid for expressing CreS<sub>N27RtoA</sub> (all Arg mutated to Ala in residues 1-27) | This study |
| pHis17-CreS<sub>Rod</sub> | pHis17-based plasmid for expressing CreS<sub>Rod</sub> (residues 80-444) | This study |
| pFE127 | pHis17-based plasmid for expressing CreS-His (CreS<sub>wt</sub> followed by GSHHHHHH) | This study |
| pHis17-CreSΔ392-412-His | pHis17-based plasmid for expressing CreSΔ392-412-His (residues 392-412 deleted) | This study |
| pHis17-CreSΔ413-433-His | pHis17-based plasmid for expressing CreSΔ413-433-His (residues 413-433 deleted) | This study |
| pHis17-CreS413-433Vim-His | pHis17-based plasmid for expressing CreS<sub>413-433Vim</sub>-His (residues 413-433 replaced by the homologous region in human vimentin) | This study |
| pMECS-NB13-HA-His | pMECS-based phagemid for expressing NB13-HA-His, a CreS-specific nanobody followed an HA tag and a 6x His tag | This study (provided by VIB) |
| pET-22b-MB13-His | pET-22b based plasmid for expressing MB13-His, a CreS-specific megabody with a C-terminal a 6x His tag | This study |
| pCT133-P<sub>creS</sub>-CreS<sub>wt</sub> | pCT133 based plasmid for expressing CreS<sub>wt</sub> under its native promoter | This study |
| pCT133-P<sub>creS</sub>-CreSΔN27 | pCT133 based plasmid for expressing CreSΔN27 (aa 1-27 deleted) under its native promoter | This study |
| pCT133-P<sub>creS</sub>-CreS<sub>M443C</sub> | pCT133 based plasmid for expressing CreS<sub>M443C</sub> under its native promoter | This study |
| pCT133-P<sub>creS</sub>-CreS<sub>E43C+S74C</sub> | pCT133 based plasmid for expressing CreS<sub>E43C+S74C</sub> under its native promoter | This study |
| pCT133-P<sub>creS</sub>-CreS<sub>E57C+E63C</sub> | pCT133 based plasmid for expressing CreS<sub>E57C+E63C</sub> under its native promoter | This study |
| pCT133-P<sub>creS</sub>-CreS<sub>R419C+S34C</sub> | pCT133 based plasmid for expressing CreS<sub>R419C+S34C</sub> under its native promoter | This study |
| pCT133-P<sub>creS</sub>-CreS<sub>R419C+A37C</sub> | pCT133 based plasmid for expressing CreS<sub>R419C+A37C</sub> under its native promoter | This study |
| pCT133-P<sub>creS</sub>-CreS<sub>K369C+T131C</sub> | pCT133 based plasmid for expressing CreS<sub>K369C+T131C</sub> under its native promoter | This study |
| pCT133-P<sub>creS</sub>-CreS<sub>K369C+A134C</sub> | pCT133 based plasmid for expressing CreS<sub>K369C+A134C</sub> under its native promoter | This study |
| pCT133-P<sub>creS</sub>-CreS<sub>R384C+T120C</sub> | pCT133 based plasmid for expressing CreS<sub>R384C+T120C</sub> under its native promoter | This study |
| pCT133-P<sub>creS</sub>-CreS<sub>K296C+Q204C</sub> | pCT133 based plasmid for expressing CreS<sub>K296C+Q204C</sub> under its native promoter | This study |
| pCT133-P<sub>creS</sub>-CreS<sub>K296C+A207C</sub> | pCT133 based plasmid for expressing CreS<sub>K296C+A207C</sub> under its native promoter | This study |
| pCT133-P<sub>creS</sub>-CreS<sub>A282C+D221C</sub> | pCT133 based plasmid for expressing CreS<sub>A282C+D221C</sub> under its native promoter | This study |
| pCT133-P<sub>creS</sub>-CreS<sub>Q204C</sub> | pCT133 based plasmid for expressing CreS<sub>Q204C</sub> under its native promoter | This study |
| pCT133-P<sub>creS</sub>-CreS<sub>K296C</sub> | pCT133 based plasmid for expressing CreS<sub>K296C</sub> under its native promoter | This study |
| pCT133-P<sub>creS</sub>-CreS<sub>T131C</sub> | pCT133 based plasmid for expressing CreS<sub>T131C</sub> under its native promoter | This study |
| pCT133-P<sub>creS</sub>-CreS<sub>K369C</sub> | pCT133 based plasmid for expressing CreS<sub>K369C</sub> under its native promoter | This study |
| pCT155-P<sub>creS</sub>-CreS<sub>WT</sub> | pCT155 based plasmid for expressing CreS<sub>wt</sub> under its native promoter | This study |

**Table S2.**
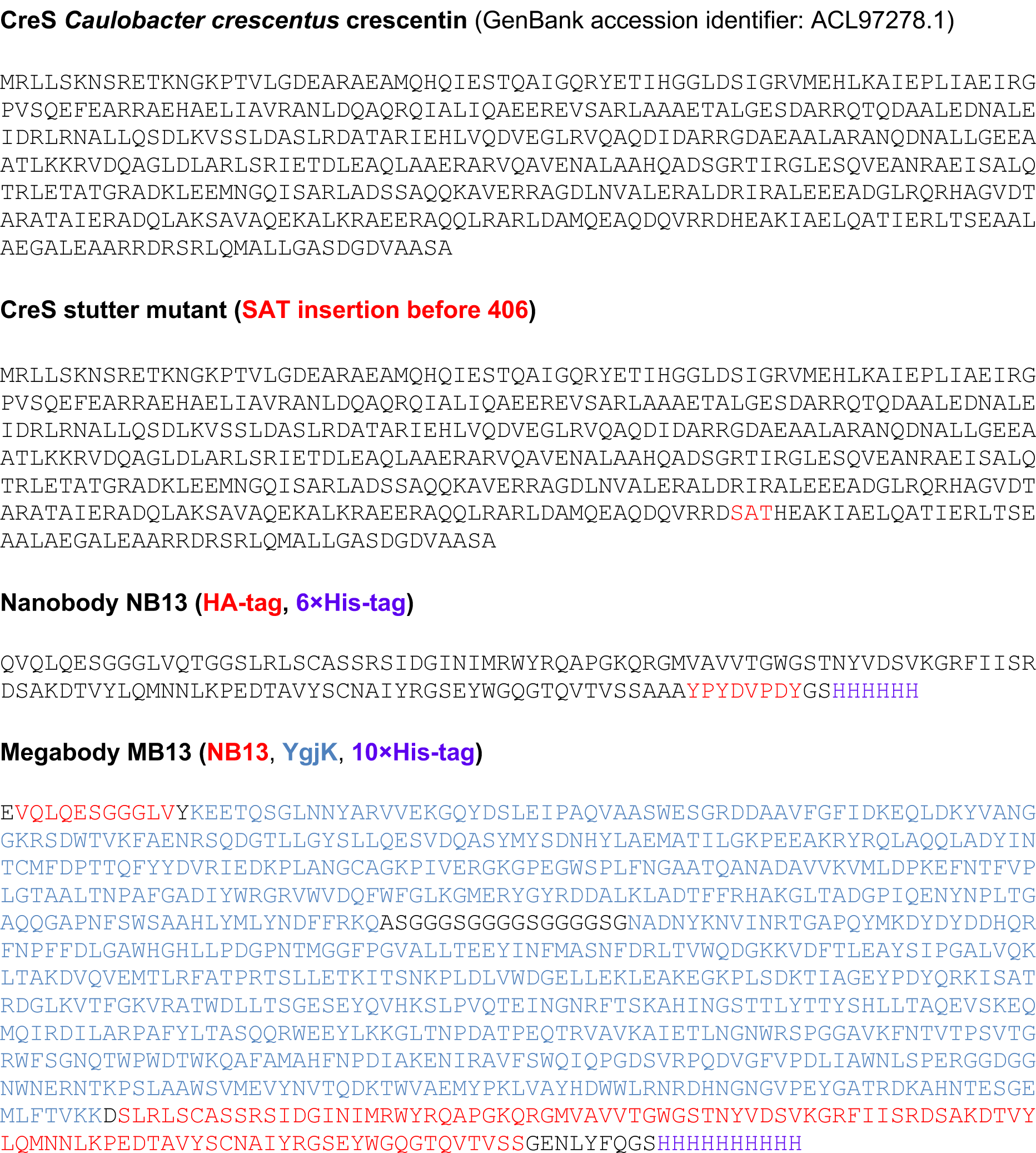
**Amino acid sequences of proteins used in this study**.

**Table S3.**
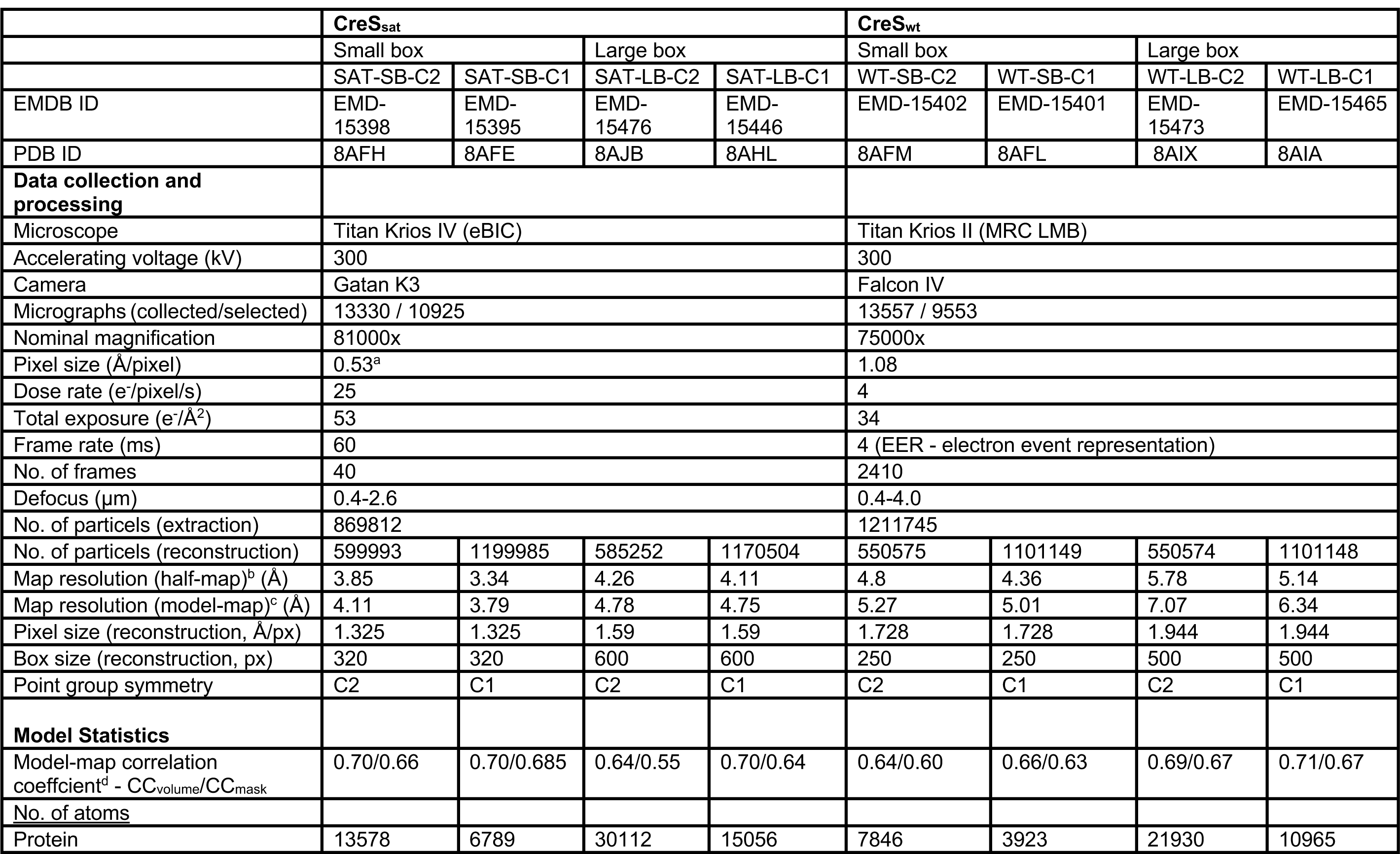
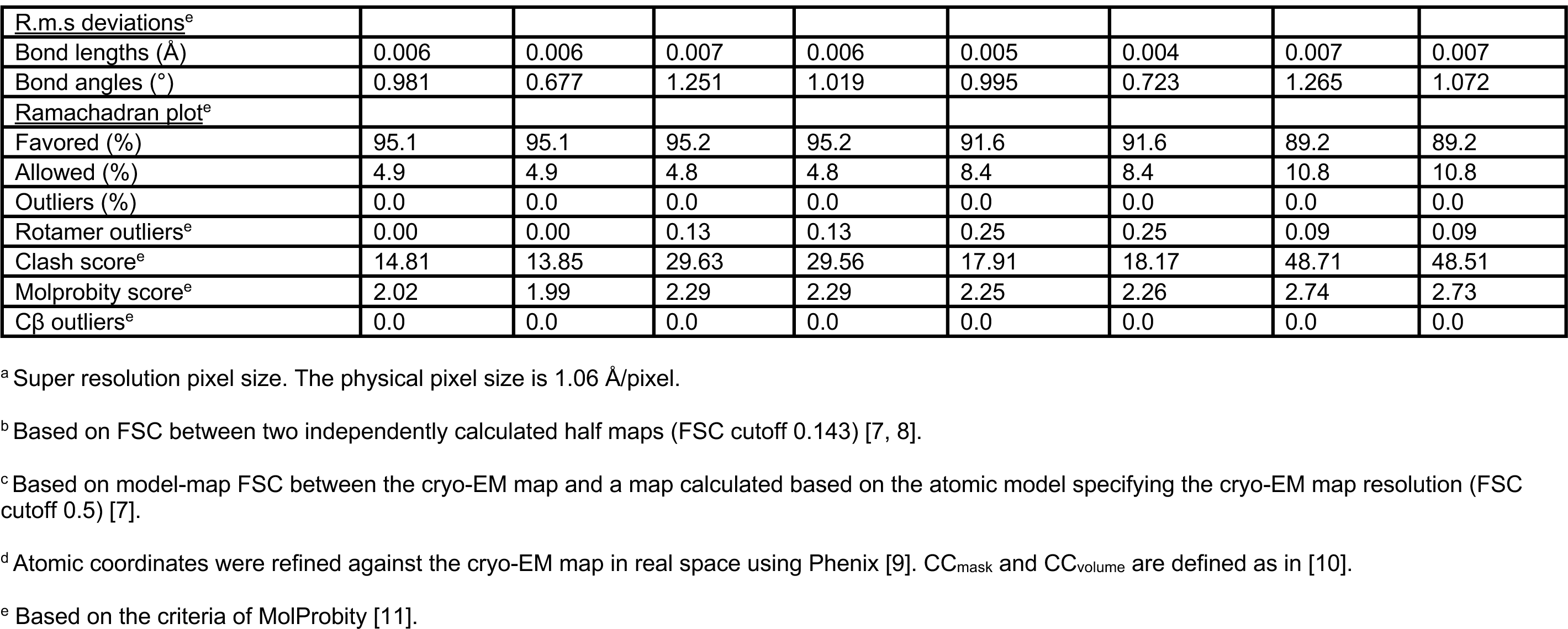
Cryo-EM data collection and processing statistics.

**Table S4.** Comparison of residue-residue contacts between CreS_sat_ and CreS_wt_.

| Region | Type | Residue A<sup>a</sup> | Residue B<sup>a</sup> | CreS<sub>wt</sub><sup>b</sup> (Å) | CreS<sub>sat</sub><sup>b</sup> (Å) |
| --- | --- | --- | --- | --- | --- |
| Middle | Inter-dimer, lateral | K369 | T131 | 8 (9) | 8 (9) |
| Middle | Intra-dimer | K369 | K369 | 17 (15) | 16 (14) |
| Middle | Intra-dimer | T131 | T131 | 16 (15) | 15 (14) |
| Middle | Inter-dimer, lateral | K369 | A134 | 7 (10) | 8 (10) |
| Middle | Inter-dimer, lateral | R384 | T120 | 8 (10) | 8 (9) |
| Crossover | Inter-dimer, lateral | K296 | Q204 | 11 (11) | 10 (11) |
| Crossover | Intra-dimer | K296 | K296 | N/A<sup>c</sup> | 16 (14) |
| Crossover | Intra-dimer | Q204 | Q204 | 17 (15) | 15 (14) |
| Crossover | Inter-dimer, lateral | K296 | A207 | 8 (10) | 7 (9) |
| Crossover | Inter-dimer, lateral | A282 | D221 | N/A<sup>c</sup> | 6 (6) |
| NN | Inter-dimer, longitudinal | E43 | S74 | 12 (13) | 7 (7) |
| NN | Inter-dimer, longitudinal | E57 | E63 | 12 (14) | 8 (9) |
| NN | Intra-monomer | E57 | E63 | 11 (10) | 12 (11) |
| CC | Inter-dimer, longitudinal | M443 (446) | M443 (446) | N/A<sup>c</sup> | 12 (10) |
| CC | Intra-dimer | M443 (446) | M443 (446) | 18 (16) | 18 (17) |
| NC | Inter-dimer, longitudinal | R419 (422) | S34 | N/A<sup>c</sup> | 12 (12) |
| NC | Inter-dimer, longitudinal | R419 (422) | A37 | N/A<sup>c</sup> | 10 (9) |

<sup>a</sup> Based on CreS<sub>wt</sub> numbering. Values in parentheses represent residue number based on CreS<sub>sat</sub> numbering.
<sup>b</sup> Cβ -Cβ distance between two residues. Values in parentheses represent the Ca-Ca distance. For a given pair, the distance shown is the shortest among all equivalent pairs related by pseudo symmetry.
<sup>c</sup> At least one of the two residues is absent in the atomic model due to disorder or an ambiguity in model building.

